# Peroxisome counteracts oxidative stresses by suppressing catalase import via Pex14 phosphorylation

**DOI:** 10.1101/2020.05.01.072132

**Authors:** Kanji Okumoto, Mahmoud El Shermely, Masanao Natsui, Hidetaka Kosako, Ryuichi Natsuyama, Toshihiro Marutani, Yukio Fujiki

## Abstract

Most of peroxisomal matrix proteins including a hydrogen peroxide (H_2_O_2_)-decomposing enzyme, catalase, are imported in a peroxisome-targeting signal type-1 (PTS1)-dependent manner. However, little is known about regulation of the membrane-bound protein import machinery. Here, we report that Pex14, a central component of the protein translocation complex in peroxisomal membrane, is phosphorylated in response to oxidative stresses such as H_2_O_2_ in mammalian cells. H_2_O_2_-induced phosphorylation of Pex14 at Ser232 suppresses peroxisomal import of catalase *in vivo* and selectively decreases formation of PTS1 receptor Pex5-mediated ternary complex of Pex14 with catalase *in vitro*. Phosphomimetic Pex14-S232D mutant elevates the level of cytosolic catalase, but not canonical PTS1-proteins, conferring higher cell resistance to H_2_O_2_. We thus suggest that H_2_O_2_-induced phosphorylation of Pex14 spatiotemporally regulates peroxisomal import of catalase, functioning in counteracting action against oxidative stress by the increase of cytosolic catalase.

## Introduction

Peroxisome, an essential intracellular organelle, functions in various essential metabolism including β-oxidation of very long chain fatty acids, and the synthesis of ether phospholipids (Waterham et al., 2016). Peroxisome contains a number of oxidases that generate hydrogen peroxide (H_2_O_2_) and catalase that decomposes H_2_O_2_ and potentially regulates reactive oxygen species (ROS) in the cell (Schrader and Fahimi, 2006). Peroxisomal functions rely on the tightly and spatiotemporally regulated import of the enzyme proteins responsible for respective reactions. Two topogenic signals are identified in the majority of peroxisomal matrix proteins: peroxisome targeting signal type-1 (PTS1) is a C-terminal tripeptide sequence SKL and its derivatives (Gould et al., 1987; Miura et al., 1992) and PTS2 is an N-terminal cleavable nonapeptide presequence (Osumi et al., 1991; Swinkels et al., 1991). Of 14 peroxisome assembly factors called peroxins in mammals, Pex14 is a peroxisomal membrane peroxin playing a central role in the import of both PTS1- and PTS2-proteins (reviewed in (Fujiki et al., 2014; Platta et al., 2016). PTS1 receptor Pex5 recognizes newly synthesized PTS1-proteins in the cytosol. Pex14 acts as an initial target of the Pex5-PTS1-protein complex on peroxisomal membrane. By associating of Pex5 with the import machinery complexes in peroxisome membrane comprising Pex14, Pex13, and RING peroxins Pex2, Pex10 and Pex12, Pex5 transports its cargo proteins into the matrix, and then shuttles back to the cytosol (reviewed in Fujiki et al., 2014; Liu et al., 2012; Platta et al., 2016).

In mammals, catalase encoded by a single gene is a tetrameric heme-containing enzyme harboring an atypical PTS1, KANL, at the C-terminus (Purdue and Lazarow, 1996). Similar to typical PTS1 proteins, catalase is mainly localized to peroxisomes by Pex5-mediated import pathway (Otera and Fujiki, 2012). Catalase forms fully active tetrameric conformation in the cytosol as noted in peroxisome-deficient fibroblasts (Middelkoop et al., 1993). Increased level of catalase is observed in the cytosol in aged human skin fibroblasts (Legakis et al., 2002). Furthermore, we recently reported that a proapoptotic protein BAK partially localizes to peroxisomes in mammalian cells and is involved in the release of catalase from peroxisomes (Fujiki et al., 2017; Hosoi et al., 2017). Although these findings suggest physiological importance of cytosolic catalase, molecular mechanisms underlying the regulation in translocation of peroxisomal matrix proteins remain largely unknown.

Posttranslational modification of protein regulates various functions of the cell in a fast, dynamic, and reversible fashion upon response to the changes in cellular demands and environmental conditions. Indeed, ubiquitination of Pex5 at a conserved cysteine residue is essential for its export from peroxisomes to the cytosol and peroxisomal matrix protein import (Carvalho et al., 2007; Okumoto et al., 2011). The cysteine residue at the position 11 of Pex5 is shown to be redox-sensitive, thereby Pex5-mediated PTS1 protein import can be regulated in the response to oxidative stress (Apanasets et al., 2014; Walton et al., 2017). As for another major posttranslational modification, i.e. phosphorylation, a large number of phosphorylation sites have been identified in various peroxisomal proteins by phosphoproteomic analysis in the yeast *Saccharomyces cerevisiae*, mouse, and humans (Oeljeklaus et al., 2016). Of these, phosphorylation of Pex14 is reported in the yeast *Hansenula polymorpha* (Komori et al., 1999; Tanaka et al., 2013) and *Pichia pastoris* (Farre et al., 2008; Johnson et al., 2001), but the biological importance and function remain unclear. In mammalian cells, mitogen-activated protein kinase (MAPK) pathways are shown to be activated in response to various oxidative stresses including ROS in the regulation of various cellular processes (Ray et al., 2012). Peroxisome is a H_2_O_2_-generating and -consuming organelle (Fransen et al., 2012), thus these findings suggest potential roles of ROS-dependent protein phosphorylation in regulating peroxisomal functions. It was reported that ataxia-telangiectasia mutated (ATM) kinase activated by ROS phosphorylates and subsequently ubiquitinates Pex5, thereby giving rise to degradation of peroxisomes, termed pexophagy (Zhang et al., 2015). However, how ROS plays a role in peroxisomal protein import remains undefined in any species.

Here, we address H_2_O_2_-dependent phosphorylation of mammalian Pex14. Phosphorylated Pex14 suppresses peroxisomal import of catalase, thereby functioning as an anti-oxidative stress response by elevating the level of catalase in the cytosol.

## Results

### Phosphorylation of Pex14 in mammalian cells

To investigate whether Pex14 is phosphorylated in mammalian cells, lysates of various mouse tissues were analyzed by electrophoresis using conventional polyacrylamide gel (SDS-PAGE) and the one containing Phos-tag (hereafter described as Phos-tag PAGE). In Phos-tag PAGE, phosphorylated proteins can be distinguished as slower-migrating bands from the corresponding non-phosphorylated form (Kinoshita et al., 2006). We found that in Phos-tag PAGE, a Pex14 band with slower migration was readily discernible by immunoblotting in the lysates of mouse testis and liver (Fig. 1A, upper panel, lanes 1 and 3, solid arrowhead) in addition to a similar level of unmodified Pex14 in both organs (open arrowhead). Pex14 was detected as a single band in conventional SDS-PAGE (Fig. 1A, middle panel). The retarded-mobility form of Pex14 completely disappeared upon treatment with **λ**-protein phosphatase (Fig. 1A, lanes 2 and 4), suggesting that Pex14 was partially phosphorylated in mammalian tissues, as observed in yeast (Johnson et al., 2001; Komori et al., 1999). Further Phos-tag PAGE analysis showed that Pex14 in mouse tissues examined was phosphorylated at varying levels (Supplementary Fig. 1A), where the relatively higher phosphorylation was detected in liver and heart (Supplementary Fig. 1A, lanes 2 and 5). Similar phosphorylation profile of Pex14 was observed in rat hepatoma Fao cells under normal culture condition (Fig. 1B, upper panel). Notably, we found that treatment with hydrogen peroxide (H_2_O_2_) increased the slower-migrating band of Pex14 with an additional lower mobility band in Phos-tag PAGE (Fig. 1B, upper panel). Treatment of Fao and CHO-K1 cells with another oxidative agent, diethyldithiocarbamate (DDC), induced nearly complete shifting of Pex14 from the unmodified form to two slower-migrating bands (Fig. 1B, upper panel). H_2_O_2_- and DDC-dependent mobility shift of Pex14 in Phos-tag PAGE was likewise observed in CHO-K1 cells, to a similar extent between two oxidative agents (Fig. 1B, upper panel). In contrast, as a negative control, neither slower-migrating bands nor unmodified Pex14 were discernible in a *PEX14*-deficient (*pex14*) CHO mutant, ZP161 (Shimizu et al., 1999) (Fig. 1B, upper panel).

**Figure 1.**
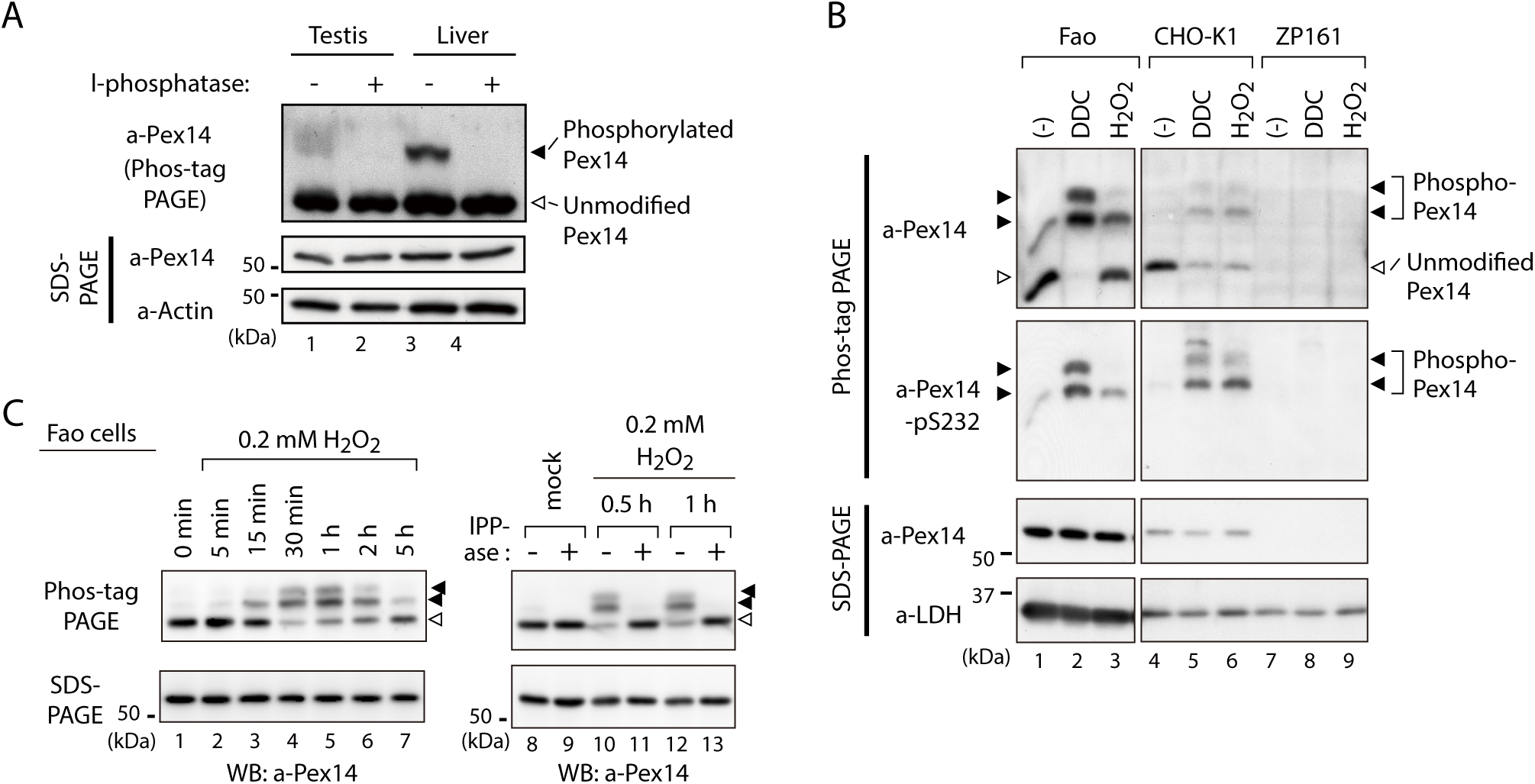
Pex14 is phosphorylated *in vivo*. (A) Lysates of testis and liver (20 µg each) from an 8-week-old male mouse were incubated with vehicle (-, lanes 1 and 3) and 400 unit λ-protein phosphatase (+, lanes 2 and 4). Samples were separated by Phos-tag PAGE (top panel) and SDS-PAGE (middle and bottom panels) and analyzed by immunoblotting with antibodies to Pex14 and actin, a loading control. Solid and open arrowheads indicate phosphorylated and unmodified Pex14, respectively. (B) Phosphorylation of Pex14 upon treatment with oxidative agents. Fao, CHO-K1, and a *PEX14*-deficient (*pex14*) CHO mutant ZP161 cells (4 × 10^5^ cells each) were treated for 30 min with vehicle (-), 100 μM diethyldithiocarbamate (DDC), and 1 mM hydrogen peroxide (H_2_O_2_). Cell lysates were analyzed as in A with antibodies to Pex14, phosphorylated Pex14 at Ser232 (Pex14-pS232), and lactate dehydrogenase (LDH). Open and solid arrowheads indicate unmodified and phosphorylated Pex14, respectively. Note that antibody to phsopho-Pex14 at Ser232 specifically recognized slower-migrating bands of Pex14 in Phos-tag PAGE. (C) *Left*, Time course of Pex14 phosphorylation upon H_2_O_2_ treatment. Fao cells were treated with 0.2 mM H_2_O_2_ as in B for indicated time periods. *Right*, λ-protein phosphatase treatment of phosphorylated Pex14. After the treatment with mock or 0.2 mM H_2_O_2_ for 0.5 h and 1 h, Fao cells were incubated with vehicle (-) and λ-phosphatase (+) as in A. The cell lysates were analyzed by Phos-tag PAGE (upper panels), SDS-PAGE (lower panels), and immunoblotting with anti-Pex14 antibody.

In Phos-tag PAGE using Fao cells, lower-migrating bands of Pex14 emerged at 15-min cell culture with 0.2 mM H_2_O_2_, peaked at 30 min to 1 h, and gradually decreased to a basal level within 5 h, while the amount of Pex14 was not altered in SDS-PAGE during the course of experiments (Fig. 1C, left panels). The H_2_O_2_-induced Pex14 bands with lower mobility were sensitive to **λ**-protein phosphatase treatment and converged to the unmodified form (Fig. 1C, right panels). These results strongly suggested that mammalian Pex14 is phosphorylated *in vivo* and that oxidative stress such as H_2_O_2_-treatment transiently enhances the phosphorylation of Pex14. We further investigated H_2_O_2_-stimulated phosphorylation of Pex14 and its functional consequence.

### Phosphorylation of Pex14 at Ser232, Ser247, and Ser252 is induced upon H_2_O_2_-treatment

Pex14 is a peroxisomal membrane protein containing a putative transmembrane segment and a coiled-coil domain (Fig. 1A, upper diagram) (Shimizu et al., 1999; Will et al., 1999). Phos-tag PAGE analysis suggested that phosphorylation of Pex14 was highly induced at several sites upon H_2_O_2_ treatment of cells. It is known that various oxidative stresses activate MAPK signaling pathways (Gaestel, 2006; Ray et al., 2012). To identify H_2_O_2_-inducible phosphorylation sites of Pex14, we expressed Pex14 mutants harboring alanine-substitutions for the potential phosphorylation residues that match with the consensus MAPK target sequence, Ser/Thr-Pro motif (Gaestel, 2006). When wild-type (WT) His-tagged rat Pex14 (His-Pex14) was expressed at a low level in Fao cells, H_2_O_2_-induced His-Pex14 phosphorylation was detected in Phos-tag PAGE (Fig. 2A), consistent with the case of endogenous Pex14 (Fig. 1, B and C). In verifying various Ala-mutants of His-Pex14, a substitution of Ser232 to Ala (S232A) eliminated two slower-migrating bands of His-Pex14 (Fig. 2A). We raised an antibody that specifically recognized the phosphorylated Ser232 of Pex14 (Supplementary Fig. 1B) and demonstrated H_2_O_2_-induced phosphorylation of endogenous Pex14 at Ser232 in Fao and CHO-K1 cells (Fig. 1B). After H_2_O_2_-treatment of cells, His-Pex14-S232A still represented as slower-migrating fuzzy bands in Phos-tag PAGE (Fig. 2A), suggesting that Ser232 is a major phosphorylation site of Pex14 and several other minor sites are present.

**Figure 2.**
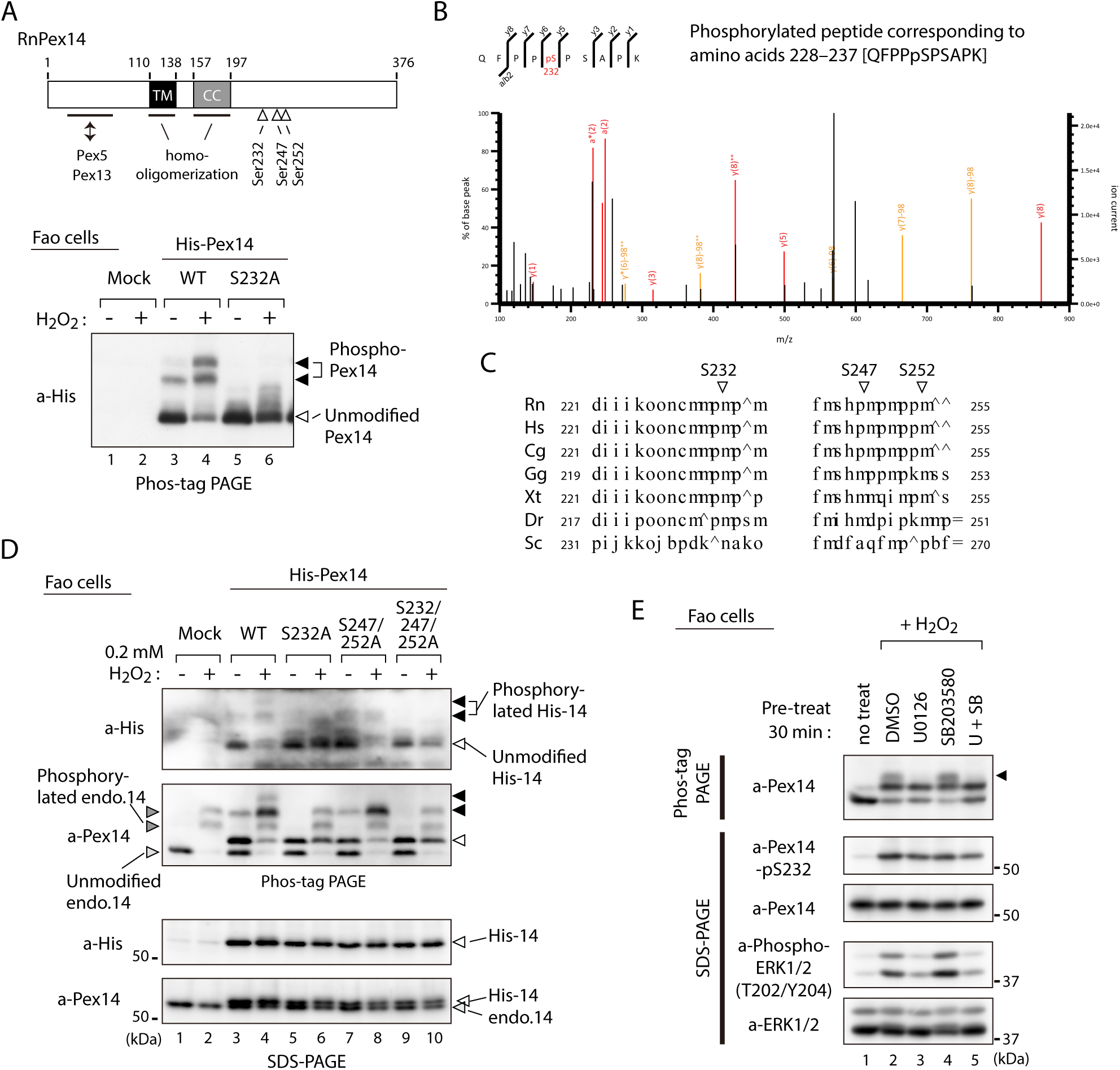
Hydrogen peroxide induces phosphorylation of Pex14 at three distinct serine residues in Fao cells. (A) *Upper*, a schematic view of domain structure of rat Pex14. Solid box, putative transmembrane (TM) domain; gray box, coiled-coil (CC) domain. *Lower*, Fao cells (4 × 10^5^ cells) were transiently transfected with plasmids encoding wild-type His-*Rn*Pex14 (WT) and S232A mutant harboring substitution at Ser232 to Ala, and a mock plasmid (mock). At 24 h after transfection, cells were treated for 30 min with vehicle (-) and 1 mM H_2_O_2_ (+) and the cell lysates were analyzed by Phos-tag PAGE and immunoblotting with anti-His antibody. Open and solid arrowheads indicate unmodified and phosphorylated forms of His-Pex14, respectively. (B) Mass spectrometric analysis of phosphorylated Pex14 induced by H_2_O_2_-treatment. Endogenous Pex14 in Fao cells (8 × 10^6^ cells) treated for 30 min with vehicle or 0.2 mM H_2_O_2_ was immunoprecipitated with anti-Pex14 antibody and subjected to LC-MS/MS analysis. Fragment spectrum of a phosphorylated peptide corresponding to amino acids 228-237 [QFPPpSPSAPK] showed phosphorylation of endogenous Pex14 at Ser232 (pS232) upon H_2_O_2_ treatment. (C) Multiple amino-acid sequence alignment of Pex14 neighboring Ser232, Ser247, and Ser252 of rat Pex14. (D) Fao cells transiently expressing wild-type His-Pex14 (WT) and the variants with indicated mutations were treated with 0.2 mM H_2_O_2_ for 30 min as in A and analyzed as in Fig. 1A with antibodies to His and Pex14. Open and solid arrowheads were as in A. (E) Fao cells (4 × 10^5^ cells) pre-incubated for 30 min with vehicle (DMSO), 10 μM U0126, 10 μM SB203580, and 10 μM U0126 plus SB203580 were further treated with 0.2 mM H_2_O_2_ for 30 min. Cell lysates were analyzed as in Fig. 1A by immunoblotting with indicated antibodies.

To determine the H_2_O_2_-induced phosphorylation sites of endogenous Pex14, Pex14 was immunoprecipitated with anti-Pex14 antibody from vehicle- or H_2_O_2_-treated Fao cells, and tryptic peptides were subjected to liquid chromatography-tandem mass spectrometry (LC-MS/MS) analysis. Four phosphorylated peptides corresponding to amino acids at alignment positions 228–237 (QFPPpSPSAPK) (Fig. 2B), 2-25 [Ap(SS)EQAEQPNQPSSSPGSENVVPR], 238–278 (IPSWQIPVKp(SPS)PSSPAAVNHHSSSDISPVSNESPSSSPGK), and 247-278 (SPSPp(SS)PAAVNHHSSSDISPVSNESPSSSPGK) (Supplementary Fig. 1C) were identified in H_2_O_2_-treated Fao cells, indicating phosphorylation at Ser3 or Ser4, Ser232, Ser247 or Ser249, and Ser251 or Ser252 in Pex14. Label-free precursor ion quantification showed that Pex14 phosphorylation at Ser232, Ser247 or Ser249, and Ser251 or Ser252 increased by 21.7-, 104.7-, and 4.0-fold upon H_2_O_2_ treatment, respectively (Supplementary Fig. 1D), suggesting that Ser232, Ser247 or Ser249, and Ser251 or Ser252 were H_2_O_2_-induced phosphorylation sites of Pex14. These Ser residues except for Ser251 in rat Pex14 reside in a consensus Ser-Pro sequence of MAPKs target (Gaestel, 2006) (Fig. 2C). Ser232 of Pex14 is conserved in vertebrates, while Ser247 and Ser252 are relatively less conserved (Fig. 2C). In contrast, H_2_O_2_ treatment gave rise to a small change in phosphorylation at Ser3 or Ser4 (2.3-fold increase) (Supplementary Fig. 1D). H_2_O_2_-induced phosphorylation of Pex14 at Ser232 was also detected in several other cultured cell lines including human HepG2, HuH7, and HeLa cells, rat RCR1 cells, and mouse embryonic fibroblasts (MEF) (Supplementary Fig. 1E), suggesting the highly conserved oxidative stress-inducible phosphorylation of Pex14 in mammals.

Next, three Ser residues, Ser232, Ser247, and Ser252, located in the C-terminal region of Pex14 (Fig. 1A, upper diagram) were serially substituted to Ala to assess respective phosphorylation upon H_2_O_2_-treatment. His-Pex14-WT showed two phosphorylated bands in Phos-tag PAGE of H_2_O_2_-treated Fao cells and S232A mutation eliminated both bands (Fig. 2D, lower panel of Phos-tag PAGE, lanes 4 and 6, solid arrowheads). In contrast, only the phosphorylated band with slower migration disappeared in the S247A/S252A double mutant (Fig. 2D, lane 8), suggesting the phosphorylation at Ser247 and/or Ser257. Phosphorylated Pex14 was undetectable in the S232A/S247A/S252A triple mutant (Fig. 2D, lane 10), consistent with the LC-MS/MS analysis (Fig. 2B). Interestingly, H_2_O_2_-induced phosphorylated band of endogenous Pex14 with slower migration was specifically eliminated by pre-treatment with ERK1/2 inhibitor U0126, but not with p38 inhibitor SB203580 (Fig. 2E, top panel, lanes 3 and 4, upper band (solid arrowhead)). Pre-treatment with both U0126 and SB203580 appeared to slightly reduce the level of phospho-Ser232 in Pex14 (Fig. 2E, second upper panel). Taken together, these results suggest that Pex14 phosphorylation at Ser232 is primarily induced upon H_2_O_2_ treatment and that Ser247 and Ser252 are phosphorylated in an ERK-mediated manner.

### Phosphorylation of Pex14 suppresses peroxisomal import of catalase, not PTS1 proteins

We next investigated whether phosphorylation of Pex14 is involved in regulation of peroxisomal import of matrix proteins. In CHO-K1 and CHO *pex14* mutant ZP161 (Shimizu et al., 1999), exogenous expression of Pex14 under a strong CMV promoter resulted in its phosphorylation without cell-treatment with H_2_O_2_ (Supplementary Fig. 2A). By introducing a modified CMV promoter lacking the enhancer region (Okatsu et al., 2012), we expressed Pex14 in ZP161 at a lower level including a phosphorylated Pex14 (Supplementary Figs. 1B and 2A). With this weaker promoter, a series of Pex14 mutants with Ser-to-Ala (as phosphorylation-defective mutants) or Ser-to-Asp substitution (as phosphomimetic mutants) were transiently expressed in a *pex14* mutant ZP161. Pex14-S232A mutant similarly restored peroxisomal import of catalase in ZP161 as wild-type Pex14, but Pex14-S232D mutant was lowered by about 50% in the restoring efficiency of the impaired catalase import (Fig. 3, A and B). In contrast, both S232A and S232D mutations showed no significant difference in restoring of PTS1 protein import (Fig. 3B). Phosphorylation-deficient Pex14 mutants with either double mutation S247A/S252A or triple mutation S232A/S247A/S252A had no effect on respective restoring activity in catalase import, whereas the phosphomimetic triple mutant S232D/S247D/S252D, not double mutant S247D/S252D, further lowered the efficacy than the single mutant S232D in the peroxisomal import of catalase (Fig. 3, A and B). In Pex14 mutants examined, peroxisomal import of PTS1 proteins was weakly, ∼ 20%, decreased in the triple mutant S232D/S247D/S252D (Fig. 3B). Together, these results suggested that Pex14 phosphorylations each at Ser232 and Ser247/Ser252 were mainly and additively involved in reducing peroxisomal import of catalase, respectively, with high specificity.

**Figure 3.**
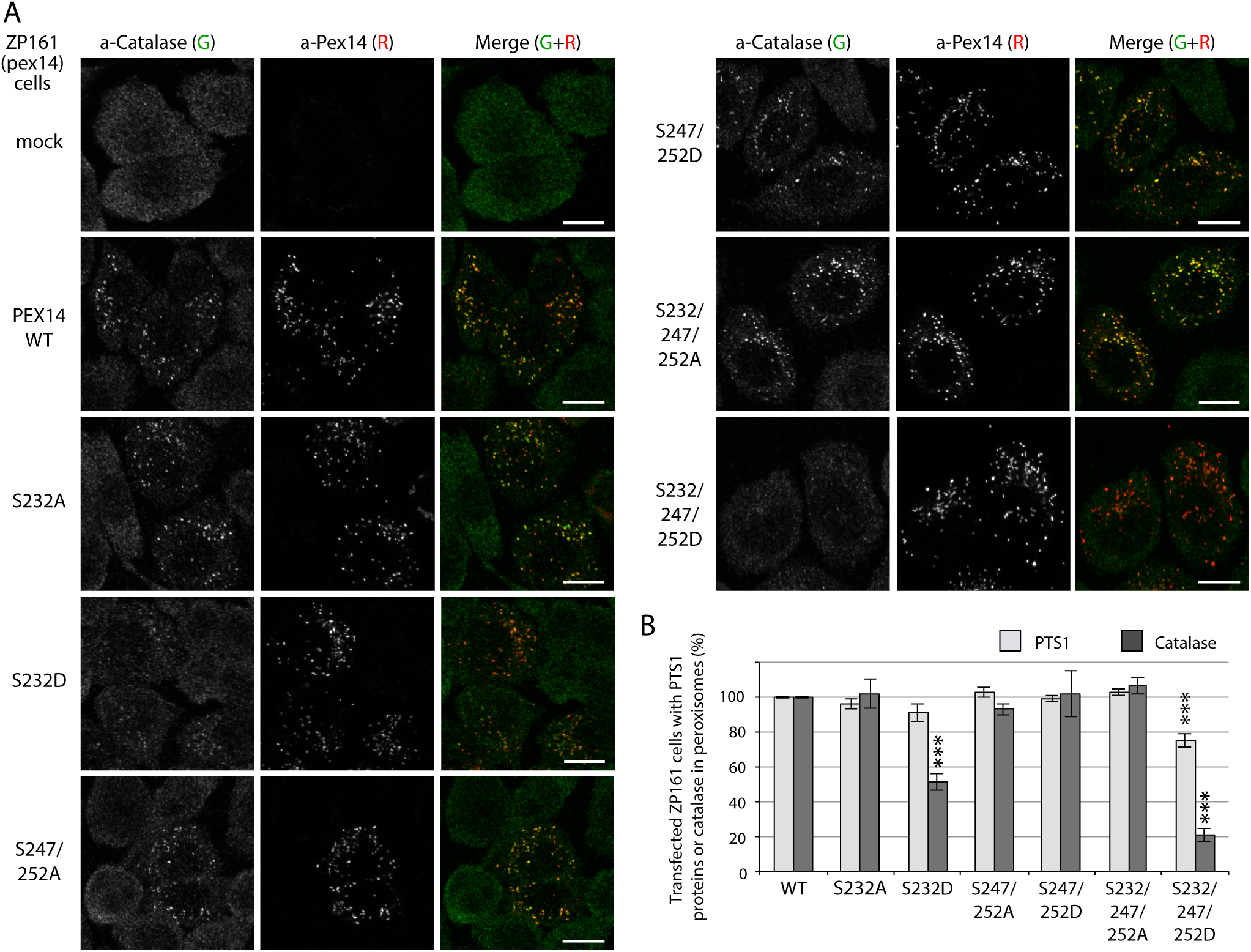
Phosphorylation of Pex14 suppresses peroxisomal import of catalase. (A) *pex14* ZP161 was transiently transfected with an empty vector (mock), or wild-type (WT) and respective Ser mutants of *FLAG-PEX14*. At 36 h after transfection, cells were immunostained with antibodies to catalase (green) and Pex14 (red). Bar, 10 μm. (B) Quantification of the data in A and those for PTS1 proteins. Percentages of the cells where PTS1 proteins (light gray) and catalase (dark gray) were mostly localized in peroxisomes in Pex14-expressing cells were represented as the means ± SD by taking those as 100% in Pex14-WT-expressing cells. Transfected cells (n > 50) were counted in three independent experiments. \*\*\**p* < 0.001; one-way ANOVA with Dunnett’s post hoc test versus cells expressing Pex14-WT.

To further examine the effect of Pex14 phosphorylation on the regulation of catalase import, we established stable cell lines of *pex14* ZP161 each expressing wild-type His-Pex14 (named WT-6) and its mutants, phosphorylation-defective Pex14-S232A (SA-13) and phosphomimetic Pex14-S232D (SD-30) (Fig. 4A). In these stable cell lines, wild-type Pex14 and the S232A and S232D mutants were expressed at similarly lower level (Fig. 4A, top panel). A PTS1 protein, 75-kDa acyl-CoA oxidase (AOx) A-chain, is imported to peroxisomal matrix and proteolytically processed to 53-kDa B-chain and 22-kDa C-chain components (Miyazawa et al., 1989). AOx B-chain was discernible at an equal level in three Pex14 variant-expressing stable cell lines of ZP161 as in CHO-K1, but not detectable in *pex14* ZP161 (Fig. 4A, upper middle panel), hence suggesting that Pex14-S232A and Pex14-S232D similarly restored peroxisomal import of AOx as the wild-type Pex14. Catalase expression level was indistinguishable between these three ZP161-stable cell lines, CHO-K1, and ZP161 (Fig. 4A, lower middle panel). In immunofluorescence microscopy, catalase in CHO-K1 was localized in Pex14-positive punctate structures, peroxisomes, whereas in ZP161 catalase was detectable in the cytosol due to no expression of Pex14 (Shimizu et al., 1999) (Fig. 4B). As in CHO-K1 cells, catalase was predominantly detected in peroxisomes in stable cell lines, WT-6 and SA-13 expressing wild-type His-Pex14 and Pex14-S232A, respectively (Fig. 4B). However, peroxisomal localization of catalase was severely lowered and cytosolic catalase was moderately elevated in the cell line SD-30 expressing Pex14-S232D (Fig. 4B), where PTS1 proteins were detectable in peroxisomes (Fig. 4C), which was consistent with efficiently processed of AOx as in CHO-K1 (Fig. 4A). We further verified intracellular localization of peroxisomal matrix proteins by subcellular fractionation analysis. A higher level of catalase was detected in cytosolic fraction (S) as compared to that in organelle fraction (P) from the SD-30 cells expressing Pex14-S232D (Fig. 4D, lanes 9 and 10, and Fig. 4E). Consistent with our earlier report (Hosoi et al., 2017), a part of catalase was present in the cytosol fraction in CHO-K1 and in WT-6 cells expressing wild-type Pex14 (Fig. 4 D, lanes 1 and 5, and Fig. 4E). Catalase was barely detectable in the cytosolic fraction from the SA-13 cells expressing Pex14-S232A (Fig. 4D, lane 7). The ratio of cytosolic (S) to total (S + P) catalase of SA-13 cells indicated significant decrease as compared to that of WT-6 cells (Fig. 4E), suggesting that Pex14-S232A more efficiently imported catalase into peroxisomes than wild-type Pex14 that was partially phosphorylated in WT-6 cells (Supplemental Fig. 1B. Precursors of PTS2 proteins, peroxisomal fatty acyl-CoA thiolase and alkyldihydroxyacetonephosphate synthase (ADAPS), are converted to their respective mature forms in peroxisomes by cleavage of the amino-terminal PTS2 presequences (de Vet et al., 1998; Honsho et al., 2008; Osumi et al., 1991; Swinkels et al., 1991). Only mature forms of thiolase and ADAPS were detected at a similar level in the organelle fractions from CHO-K1 and three ZP161-stable cell lines (Fig. 4D), demonstrating normal import of PTS2-proteins, similarly to the case of AOx import. These characteristics of peroxisomal matrix protein import in SD-30 cells were similarly observed in other two independent ZP161-stable cell lines expressing Pex14-S232D (data not shown). Collectively, these results suggested that phospho-mimic Pex14-S232D specifically reduces peroxisomal import of catalase, not PTS1- and PTS2-proteins.

**Figure 4.**
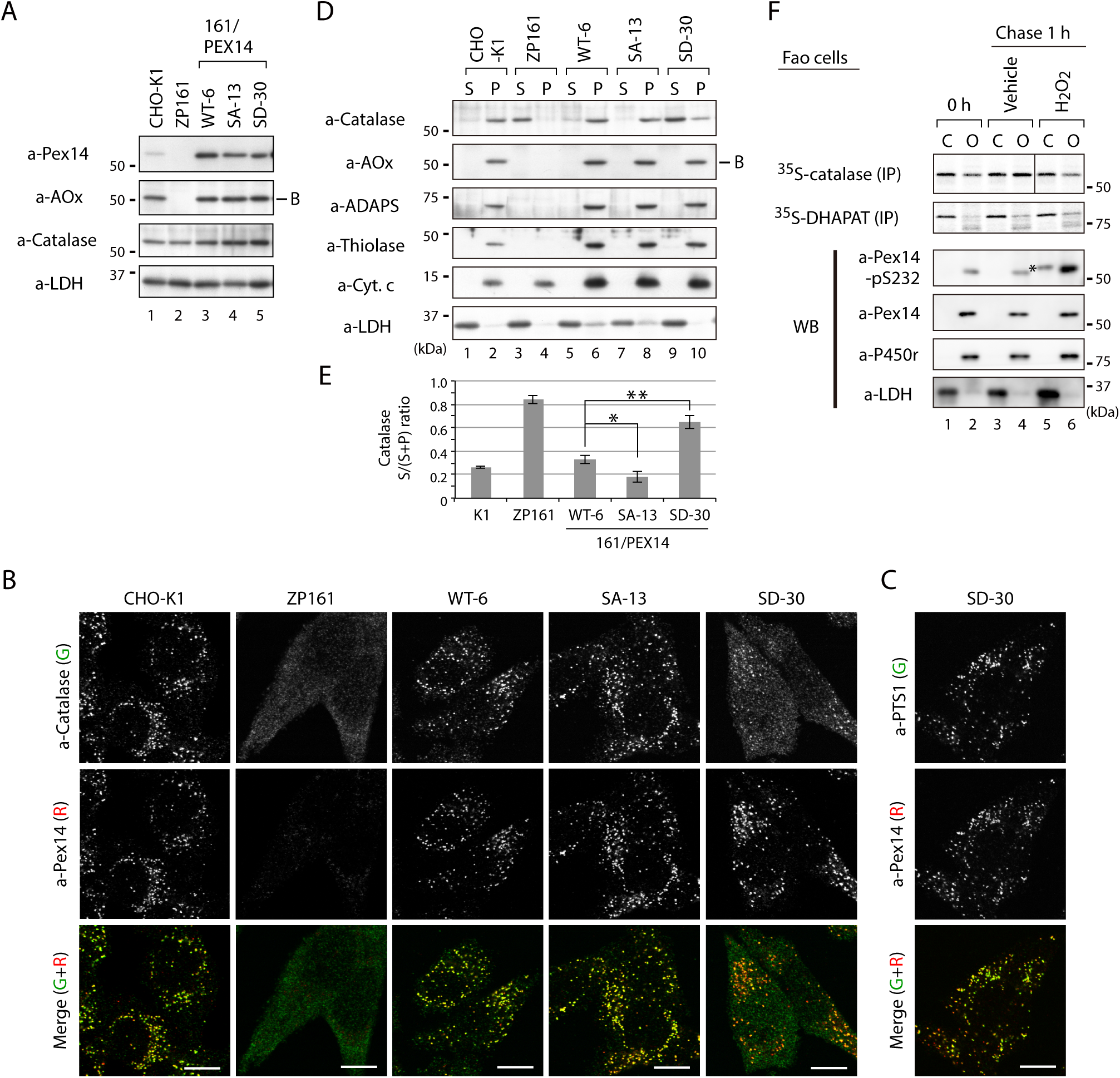
Phospho-mimic Pex14 mutant, Pex14-S232D, reduces catalase import into peroxisomes. (A) Cell lysates of CHO-K1, *pex14* ZP161, and stable cell lines of ZP161 each expressing wild-type His-Pex14 (WT-6) and its mutants, phosphorylation-deficient Pex14-S232A (SA-13) and phosphomimetic Pex14-S232D (SD-30) (4 × 10^5^ cells each) were analyzed by SDS-PAGE and immunoblotting with antibodies indicated on the left. Only the B-chain of acyl-CoA oxidase (AOx) that is generated by intraperoxisomal proteolytic processing of full-length AOx is shown. (B) Catalase was less imported into peroxisomes in Pex14-S232D-expressing cells. CHO-K1, *pex14* ZP161, and stable cell lines each expressing Pex14 variants were immunostained with antibodies to catalase (green) and Pex14 (red). Merged images were also shown. Bar, 10 μm. (C) Stable lines of *pex14* ZP161 expressing phosphomimetic Pex14-S232D (SD-30) were likewise immunostained with antibodies to PTS1 (green) and Pex14 (red). Bar, 10 μm. (D) Catalase in cytosolic fraction was increased in Pex14-S232D-expressing cells. Cells indicated at the top (8 × 10^5^ cells each) were separated into cytosolic (S) and organelle (P) fractions by permeabilization with 25 μg/mL digitonin and subsequent ultracentrifugation. Equal aliquots of respective fractions were analyzed by SDS-PAGE and immunoblotting with the indicated antibodies. AOx, a typical PTS1 protein; alkyl-dihydroxyacetonephosphate synthase (ADAPS) and 3-ketoacyl-CoA thiolase (thiolase), PTS2 proteins; Cyt. *c*, cytochrome *c*. LDH is a marker for cytosolic fraction. (E) Catalase level in cytosolic and organelle fractions assessed in D was quantified and shown as a ratio of cytosol (S) to total (S plus P). Data represent means ± SD of three independent experiments. Statistical analysis was performed by one-way ANOVA with Dunnett’s post hoc test as compared with the S/(S+P) ratio of catalase in WT-6 cells. \**p* < 0.05 and \*\**p* < 0.01. (F) Pulse-chase experiment of catalase translocation. Fao cells were labeled with ^35^S-methionine and ^35^S-cysteine for 1 h and were chased for 1 h in the presence of vehicle or 0.2 mM H_2_O_2_. Cells were fractionated into the cytosol (C) and organelle (O) fractions as described in the STAR Methods. Equal aliquots of respective fractions were solubilized and subjected to immunoprecipitation with antibodies to catalase and DHAPAT. ^35^S-labeled catalase was analyzed by SDS-PAGE and detected by autoradiography (two upper panels). Equal aliquots of the cytosol and organelle fractions were analyzed by SDS-PAGE and immunoblotting using indicated antibodies. P450r, an ER membrane protein, cytochrome P450 reductase; LDH, a cytosolic protein. *, a nonspecific band.

To investigate peroxisomal import of catalase upon the H_2_O_2_ treatment *in vivo*, Fao cells were pulse-labeled with ^35^S-labeled methionine and cysteine for 1 h and fractionated into cytosolic and organelle fractions. Immunoprecipitated ^35^S-catalase was more detected in the cytosolic fraction than that in organelle fraction at the start of chase (Fig. 4F, lanes 1 and 2). At 1-h chase in normal condition, approximately a half of ^35^S-catalase was detected in the organelle fraction, indicating peroxisomal import of newly synthesized catalase (Fig. 4F, lanes 3 and 4). On the other hands, in the presence of H_2_O_2_, the amount of ^35^S-catalase in organelle fraction after 1-h chase was almost the same level as that at 0-h chase, where Pex14 was highly phosphorylated (Fig. 4F, lanes 5 and 6). By contrast, ^35^S-labeled dihydroxyacetonephosphate acyltransferase (DHAPAT), an enzyme with typical PTS1, was increased in organelle fractions during 1-h chase in the absence or presence of H_2_O_2_ (Fig. 4F). These results strongly suggest that peroxisomal import of newly synthesized endogenous catalase is selectively suppressed by induction of Pex14 phosphorylation during the cell-exposure to H_2_O_2_.

Less efficient import of PTS1 proteins into peroxisomes and accumulation of Pex5 in peroxisome membrane are observed in human fibroblast cells at late passage of cell culture. where intracellular ROS is elevated (Legakis et al., 2002). To assess the effect of H_2_O_2_ treatment on Pex5 recycling between peroxisomes and the cytosol, subcellular fractionation was performed under the condition that enabled to detect mono-ubiquitinated Pex5 at Cys11 (Okumoto et al., 2011). Pex5 in organelle fraction was subtly and modestly increased upon 1-h H_2_O_2_-treatment at concentration of 0.2 mM (standard setting otherwise mentioned in this study) and 0.5 mM, where both concentrations of H_2_O_2_ gave rise to phosphorylation of Pex14 at Ser232 to a similar extent (Supplementary Fig. 2B, left panel, immunoblot with anti-Pex14-S232 antibody). Mono-ubiquitinated Pex5, which was detected as a DTT-sensitive slow-migrating band of Pex5 in organelle fraction, was not altered in a ubiquitinated protein level upon H_2_O_2_ treatments (Supplementary Fig. 2B, right panel, solid arrowhead). These results suggested that at least the treatment with 0.2 mM H_2_O_2_ for a short time period did not apparently affect the Pex5 recycling, as noted in the nearly normal import of ^35^S-PTS1 protein (Fig. 4F). Collectively, upon treatment of cells with H_2_O_2_ the phosphorylation of Pex14 most likely suppresses peroxisomal import of catalase more selectively than that of PTS1 proteins.

### Phosphorylation of Pex14 selectively affects Pex5-mediated import complex formation with catalase

We next investigated molecular mechanism underlying how phosphorylation of Pex14 regulates peroxisomal import of catalase. In mammals, N-terminal region of Pex14 is shown to interact with Pex5 and Pex13 (Fig. 2A, upper diagram) (Itoh and Fujiki, 2006; Otera et al., 2000; Schliebs et al., 1999). Organelle fractions from Fao cells treated with vehicle or H_2_O_2_ were subjected to immunoprecipitation with anti-Pex14 antibody. Endogenous Pex14 was equally recovered from cells with respective treatments, where the level of phosphorylated Pex14 at Ser232 was elevated in H_2_O_2_-treated cells (Fig. 5A, two top panels, lanes 3 and 4). A higher level of Pex13 was included in the Pex14 complex from H_2_O_2_-treated cells than that from vehicle-treated cells, where Pex13 was expressed apparently at the same level (Fig. 5A, lanes 1 and 2). Pex5 was increased in the organelle fraction upon H_2_O_2_ treatment (Fig. 5A, lanes 1 and 2), but in the Pex14 immunoprecipitates a lower amount of Pex5 at an equal level between the prior to and upon the treatment with H_2_O_2_ was discernible (Fig. 5A, lanes 3 and 4). These results suggested that apparently H_2_O_2_-dependent phosphorylation of Pex14 alters its complex formation with Pex13 in peroxisomal membrane. This is consistent with the finding that Pex13 is involved in peroxisomal import of catalase by interacting with Pex5 (Otera and Fujiki, 2012).

**Figure 5.**
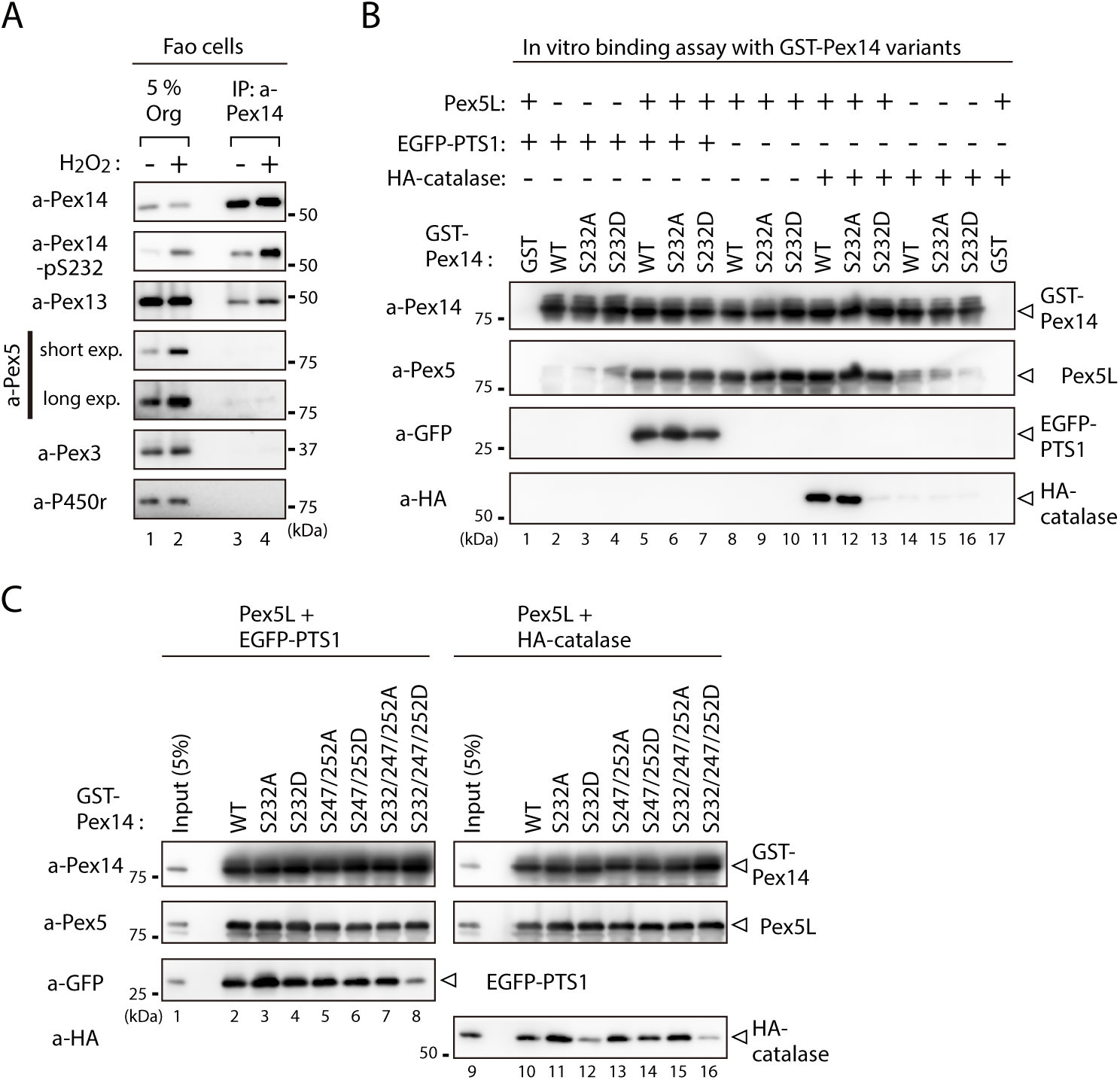
Phosphorylation of Pex14 exclusively affects Pex5-mediated complex formation with catalase. (A) Phosphorylated Pex14 forms a complex with Pex13. Organelle fractions of Fao cells (4 × 10^6^ cells each) treated for 30 min with vehicle (-) or 0.2 mM H_2_O_2_ were solubilized and subjected to immunoprecipitation with anti-Pex14 antibody. Equal-volume aliquots of immunoprecipitates (IP) and the input of organelle fractions (Org. input, 5%) were analyzed by SDS-PAGE and immunoblotting with antibodies indicated on the left. (B) *In vitro* binding assays were performed using recombinant proteins, i.e. GST-Pex14 variants, Pex5L, EGFP-PTS1, and HA-catalase. Components added to the assay mixtures, including GST in place of GST-Pex14 variants, are indicated at the top. Pex5L, EGFP-PTS1, and HA-catalase in fractions bound to GST-Pex14-conjugated glutathione-Sepharose beads were analyzed by immunoblotting with antibodies indicated on the left. (C) *In vitro* binding assays were likewise performed using GST-Pex14 variants with mutations in three distinct Ser residues as in B. Five percent input of the reaction used was also loaded.

PTS1 receptor Pex5 recognizes PTS1 cargo proteins including catalase and transports them to peroxisomes by docking on Pex14 (Fujiki et al., 2014; Platta et al., 2016). To further assess the effect of Pex14 phosphorylation in the interaction with known Pex14-binding partners, glutathione S-transferase (GST) pull-down assays were performed using recombinant proteins including a longer isoform of Pex5, Pex5L (Otera et al., 1998). In regard to Pex14-Pex5 interaction, Pex5Lwas equally detected in the fractions bound to GST-fused wild-type Pex14 (GST-Pex14-WT), GST-Pex14-S232A, and GST-Pex14-S232D, but not GST alone, suggesting that both S232A and S232D mutations have no apparent effect on direct binding of Pex14 to Pex5L (Fig. 5B, lanes 1, 8-10, 17). As shown in the earlier report (Otera and Fujiki, 2012; Otera et al., 2002), PTS1 cargoes, both EGFP-PTS1 and HA-catalase, respectively formed a ternary complex with GST-Pex14-WT via Pex5L (Fig. 5B, lanes 2, 5, 11, 14). However, GST-Pex14-S232D, not GST-Pex14-S232A, yielded almost undetectable amount of HA-catalase in the bound fraction, despite the same-level recovery of Pex5L (Fig. 5B, lanes 12 and 13). By contrast, in the presence of Pex5L, EGFP-PTS1 was detected in the fractions bound to GST-Pex14-S232D at slightly lower level, as compared to that with GST-Pex14-WT and GST-Pex14-S232A (Fig. 5B, lanes 5-7). In the absence of Pex5L, EGFP-PTS1 and HA-catalase were not detectable in the bound fractions of GST-Pex14 variants (Fig. 5B, lanes 2-4, 14-16). Essentially the same results were obtained with a shorter isoform of Pex5, Pex5S (Supplementary Fig. 3A). Together, Pex14-S232D most likely interacts with Pex5 as wild-type Pex14 but it much less efficiently forms a Pex5-mediated ternary complex with catalase. We further investigated the effect of Pex14 phosphorylation at Ser247 and Ser252 on the ternary complex formation. None of the mutations S232A and S232D, double mutations S247A/S252A and S247D/S252D, and triple mutations S232A/S247A/S252A and S232D/S247D/S252D in Pex14 altered the binding efficiency to Pex5L (Supplementary Fig. 3B). HA-catalase was similarly pulled down with GST-Pex14 harboring respective mutations, S247A/S252A, S247D/S252D, and S232A/S247A/S252A in the presence of Pex5L, as seen with GST-Pex14-WT (Fig. 5C, lanes 10, 13-15). However, a triple mutant GST-Pex14-S232D/S247D/S252D showed further decrease in the Pex5-dependent recovery of HA-catalase as compared to a single mutant GST-Pex14-S232D (Fig. 5C, lanes 12 and 16). In forming the Pex5-mediated ternary complex with EGFP-PTS1, significant decrease was observed only with the GST-Pex14-S232D/S247D/S252D (Fig. 5C, lanes 2-8). Collectively, these results suggested that phosphorylation of Pex14 at Ser232 selectively suppresses the Pex5-mediated complex formation with catalase, where the phosphorylation at Ser247 and Ser252 likely provides additive effect. Such distinct effects of Pex14 mutants on the ternary complex formation with catalase and typical PTS1 protein are in good agreement with the phenotypes of those observed *in vivo*, in cultured cells (Figs. 3 and 4).

### Phosphorylation at Ser232 of Pex14 shows higher cell resistance to hydrogen peroxide

A part of catalase is localized in the cytosol even in the wild-type CHO-K1 cells, while catalase is fully diffused to the cytosol, in peroxisome-defective mutants such as *pex14* ZP161 cells. Such cytosolic catalase is responsible for the cell resistance to exogenously added H_2_O_2_ (Hosoi et al., 2017). We next investigated whether H_2_O_2_-induced phosphorylation of Pex14 increases cytosolic catalase by suppressing its peroxisomal import in order to eliminate the cytosolic H_2_O_2_ for cell survival. At 16 h after treatment of H_2_O_2_, cell viability in *pex14* ZP161 cells was higher than in CHO-K1 cells (Fig. 6), as previously shown (Hosoi et al., 2017), and ZP161-stable cell line WT-6 expressing wild-type Pex14 (Fig. 6). A ZP161-stable cell line SD-30 expressing Pex14-S232D was more resistant to exogenous H_2_O_2_ like ZP161 than WT-6 cells, whereas a ZP161-stable cell line SA-13 expressing Pex14-S232A showed a significant decrease in the cell viability as compared with WT-6 cells (Fig. 6). Moreover, these differential sensitivities to H_2_O_2_ between ZP161-stable cell lines were completely abrogated by the addition of 3-aminotriazole, a catalase inhibitor, indicative of catalase-dependent cell viability. The level of cell resistance to exogenous H_2_O_2_ (Fig. 6) is well correlated with the amount of cytosolic catalase in respective types of cells (Fig. 4, D and E). Taken together, these results suggest that oxidative stresses such as H_2_O_2_ enhances the phosphorylation of Pex14 at Ser232, thereby suppressing peroxisomal import of catalase and concomitantly elevating the cytosolic catalase to counteract H_2_O_2_ in the cytosol for cell survival.

**Figure 6.**
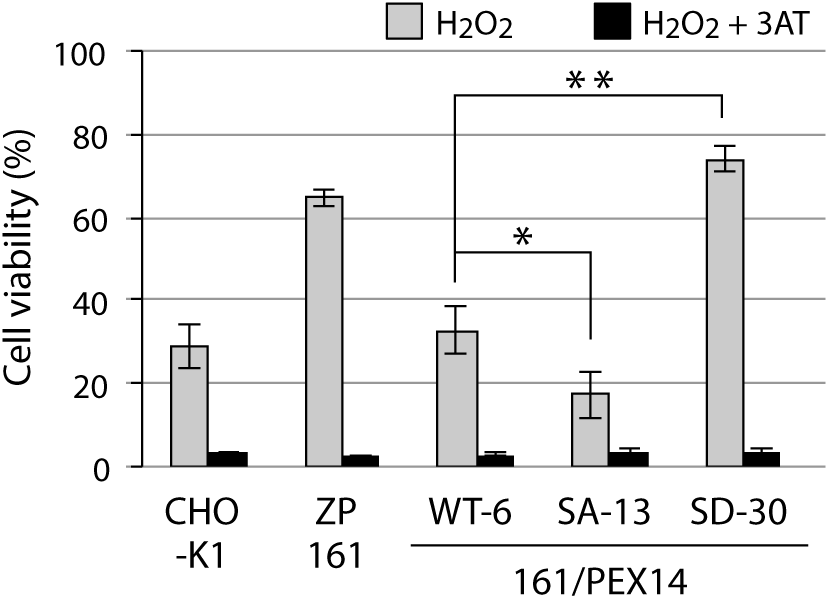
Phosphorylation at Ser232 of Pex14 is important for cell resistance to exogenous hydrogen peroxide. CHO-K1, *pex14* ZP161, and its stable cell lines expressing Pex14 variants (1 × 10^4^ cells each) were treated with 0.8 mM H_2_O_2_ in the absence (gray bars) and presence (solid bars) of 20 mM 3-aminotriazole (3AT), a catalase inhibitor. Cell viability was determined by MTS assay at 16 h after H_2_O_2_ treatment and represented as percentages relative to that of each mock-treated, H_2_O_2_-untreated cells. Data represent means ± SD of three independent experiments. \**p* < 0.05 and \*\**p* < 0.01; one-way ANOVA with Dunnett’s post hoc test versus a stable cell line of ZP161 expressing Pex14-WT.

## Discussion

Here our findings demonstrated the phosphorylation of mammalian Pex14 in response to H_2_O_2_ and assigned it as a novel regulation of peroxisomal import of catalase. We identified H_2_O_2_-induced three phosphorylation sites of Pex14, Ser232, Ser247, and Ser252 (Fig. 2B and Supplementary Fig. 1C), all of which locate in the cytosolically faced, C-terminal region of Pex14 (Fig. 2A, upper diagram). Our earlier domain mapping study suggested that the C-terminal region following the coiled-coil domain of Pex14 plays a role in peroxisomal protein import (Itoh and Fujiki, 2006). Notably, deletion of the residues at 201-367 including Ser232, not that at 261-367, abolished peroxisomal import of catalase but retained minimum import of PTS1 proteins (Itoh and Fujiki, 2006). These results support our view that C-terminal region of Pex14 has a regulatory role in peroxisomal protein import, which is modulated by phosphorylation of Ser232. Indeed, phosphorylation of Pex14 at Ser232 selectively lowers the Pex5-mediated complex formation with catalase (Fig. 5, B and C). Pex14 forms a highly ordered homo-oligomer (Itoh and Fujiki, 2006) as well as large protein complexes of protein translocation machinery (Meinecke et al., 2010). Therefore, phosphorylation at Ser232 most likely induces the conformational change *per se* and Pex14-containing protein complexes, thereby leading to the predominant suppression of catalase import into peroxisomes. Phosphorylation at Ser247 and Ser252 potentially provides additive effect on the conformation of Pex14 complex, resulting in further inhibition in the peroxisomal import of catalase as well as partial retardation in that of PTS1 proteins (Fig. 3, A and B). Upon H_2_O_2_ treatment, the level of Pex13 is increased in the immunoprecipitated Pex14 complexes containing highly phosphorylated Pex14 (Fig. 5A). Since Pex13 plays an essential role in import of catalase in mammalian cells (Otera and Fujiki, 2012), catalase import is more likely regulated in a tightly coordinated manner of the phosphorylated Pex14 with Pex13. Although phosphorylation of Pex14 at its C-terminal part is identified in several yeast species, including Thr248 and Ser258 in *H*. *polymorpha* (Komori et al., 1999; Tanaka et al., 2013) and Ser266 and Ser313 in *S*. *cerevisiae* (Albuquerque et al., 2008), their functional roles remain to be defined. Ser232 of Pex14 is conserved only in vertebrates (Fig. 2C) and the amino-acid sequences of the C-terminal region of Pex14 share lower similarity between vertebrates and yeasts, hence suggesting that functional significance of the phosphorylation of Pex14 at C-terminal region is distinct between species. Yeast and worms have cytosolic catalase in addition to peroxisomal catalase harboring PTS1-like sequence (Hartman et al., 2003), implying that the modulated intracellular distribution of catalase is beneficial to the survival of organisms during the evolution.

ROS activates various cellular signaling pathways. ATM is reported as a peroxisome-localized kinase activated by ROS, mediating Pex5 phosphorylation and induction of pexophagy (Zhang et al., 2015). An ATM inhibitor KU55933 that abrogates H_2_O_2_-indeced phosphorylation of Pex5 in HEK293 cells (Zhang et al., 2015) shows no effect on phopsphorylation of Pex14 upon the treatment with either H_2_O_2_ or DDC in Fao cells (Supplementary Fig. 3C). Therefore, H_2_O_2_ most likely induces posttranslational modification of Pex5 and Pex14 in a manner independent from ATM; the ATM-dependent phosphorylation and subsequent ubiquitination of Pex5 are distinct from the phosphorylation of Pex14 by the undefined kinase(s) reported here. Although the study with kinase inhibitors suggests the ERK-mediated phosphorylation of Pex14 at Ser247 and Ser252 (Fig. 2E), further studies may be required to identify the kinase(s) that directly phosphorylates Pex14 in an H_2_O_2_-dependent manner and its upstream signaling pathway. Moreover, Pex14 phosphorylation upon cell treatment with H_2_O_2_ is transiently induced but is gradually reverted to the unmodified form (Fig. 1C), implying the presence of phosphatase catalyzing dephosphorylation of Pex14.

We show that H_2_O_2_-induced phosphorylation of Pex14 predominately suppresses peroxisomal import of catalase more efficiently than typical PTS1 proteins. Together with the findings that H_2_O_2_ treatment transiently induces Pex14 phosphorylation (Fig. 1C) and that cell toxicity of H_2_O_2_ is more efficiently detoxified by cytosolic catalase (Fig. 6) (Hosoi et al., 2017), we propose a working model that Pex14 phosphorylation plays an essential role in the acute cell response against H_2_O_2_ challenge (Fig. 6). This is possibly a regulatory system by taking advantage of specific suppression of peroxisomal import of catalase, not that of PTS1 proteins, and temporal increase of catalase in the cytosol by phosphorylation of Pex14. However, catalase chronically residing in the cytosol compromises the redox homeostasis in peroxisomes, thereby resulting in mitochondrial dysfunction or cell senescence (Ivashchenko et al., 2011; Koepke et al., 2007; Walton et al., 2017). Given the finding that catalase catabolizes H_2_O_2_ at the highest rate without effecting on the reducing equivalent such as glutathione (Sies et al., 2017), it is reasonable to have catalase function as the first defender in acute phase upon excess H_2_O_2_ insult. Oxidative stress such as H_2_O_2_ has been suggested to affect PTS1 protein import via Pex5 modification, depending on the redox state of the conserved Cys11 residue (Apanasets et al., 2014; Walton et al., 2017). Therefore, oxidative stress intrinsically lowers the import of PTS1 proteins and catalase under the imbalance of cellular redox. Collectively, compromised import of catalase under the oxidative condition is most likely reflecting severely impaired formation of ternary complex of Pex14, Pex5, and catalase (Fig. 5, B and C) and weaker affinity of Pex5 to catalase as compared to canonical PTS1 proteins (Koepke et al., 2007; Otera and Fujiki, 2012). In *in vivo* situation, both mechanisms might simultaneously take place for the cell survival, where suppression of peroxisomal catalase import and BAK-mediated release of catalase from the peroxisomal matrix are involved (Hosoi et al., 2017).

This report demonstrates that the protein import machinery of peroxisomes plays a crucial role in the regulatory network that counteracts the exogenous oxidative stresses. There are other intracellular sources of H_2_O_2_ involving mitochondria, NADPH oxidases, and peroxisomes (Sies et al., 2017). We observed that Pex14 phosphorylation at Ser232 was induced in Fao cells upon either treatments with rotenone, a mitochondrial complex I inhibitor, or peroxisome proliferators including clofibrate and bezafibrate that are known to elevate intracellular level of H_2_O_2_ (Tada-Oikawa et al., 2003; Zhang et al., 2015) (data not shown). Thus, H_2_O_2_ generated by various stimuli in response to the change of intracellular or extracellular environments could modulate and fine-tune the intracellular localization of catalase via Pex14 phosphorylation. In mitochondrial biogenesis, phosphorylation of yeast mitochondrial outer membrane proteins, Tom20, Tom22, and Tom70, indeed regulates the import of mitochondrial proteins in a nutrient condition-dependent manner (Gerbeth et al., 2013; Schmidt et al., 2011). Mice genetically overexpressing or deleting catalase reveal that catalase is also involved in various physiological and pathological processes such as renal injury (Hwang et al., 2012) and cardiomyocyte dysfunction (Ye et al., 2004) in diabetes. Together with the biological relevance of cytosolic catalase to mitochondrial dysfunction (Ivashchenko et al., 2011) and cell senescence (Koepke et al., 2007; Walton et al., 2017), a tackling issue needs to be addressed in regards to mechanisms underlying how Pex14 phosphorylation-dependent, spatiotemporal regulation and dysregulation of catalase are linked to the oxidative-stress state or age-related disease.

## Materials and Methods

### Cell culture and DNA transfection

CHO-K1 cell, a *pex14* CHO cell mutant ZP161 (Shimizu et al., 1999), rat astrocytoma RCR1 cell, and rat hepatoma Fao cell were cultured at 37°C in Ham’s F-12 medium supplemented with 10% FBS under 5% CO_2_ and 95% air (Okumoto et al., 2011). Human cervix epitheloid carcinoma HeLa, human hepatocellular carcinoma HepG2 and HuH7 cells, and MEF cells were cultured at 37°C in DMEM (Invitrogen) supplemented with 10% FBS (Abe et al., 2018). CHO and Fao cells were transfected with DNA using Lipofectamine reagent (Invitrogen) or polyethylenimine (PEI-MAX, Polysciences) according to the manufacturer’s instructions. Stable transformants of ZP161 expressing rat Pex14 variants tagged with N-terminal hexahistidine (His-Pex14) were isolated by transfection of pcDNAZeo-D/*His-RnPEX14* variants (see below) followed by selection with Zeocin (Invitrogen), as described (Okumoto et al., 2000).

### Plasmids

Plasmids encoding rat Pex14 variants with Ser-to-Ala or Ser-to Asp substitutions were generated by an inverse PCR method (Weiner et al., 1994) with KOD-plus DNA polymerase (Toyobo) and pCMVSPORT/*His-RnPEX14* (Itoh and Fujiki, 2006) as a template. To generate plasmid for weak Pex14 expression, upstream 564 bp of the CMV promoter was deleted from pcDNA3.1/Zeo (Invitrogen) by an inverse PCR method, yielding a weaker expression vector, named pcDNAZeo-D. cDNAs encoding His-Pex14 variants in pCMVSPORT1 were ligated into the EcoRI-PstI sites in pcDNAZeo-D, generating pcDNAZeo-D vector-encoding His-Pex14 variants. Expression plasmids for GST-fused Pex14 variants were constructed by replacing the NheI-PstI fragment of wild-type *PEX14* in pGEX6P-2/*RnPEX14* (Itoh and Fujiki, 2006) with the corresponding fragment of *PEX14* variants in pCMVSPORT1 vector. To construct GST-fusion protein with catalase, the BglII–SalI fragment of *HA-Catalase* amplified by PCR from pUcD3/*HA-HsCatalase* (Otera and Fujiki, 2012) was cloned into the BamHI–SalI sites of pGEX6P-1 (GE Healthcare), thereby generating pGEX*/HA-HsCatalase*. Plasmids for GST-fusion proteins with Chinese hamster (*Cl*)Pex5S and *Cl*Pex5L (Otera et al., 2002) and EGFP-His-PTS1 (Okumoto et al., 2011) were also used. Primers used for PCR were shown in Key Resources Table.

### Antibodies and Chemicals

Antibodies used were rabbit polyclonal antibodies each to C-terminal 19-amino acid residues of Pex14 (Shimizu et al., 1999), Pex5 (Otera et al., 2000), Pex13 (Mukai and Fujiki, 2006), Pex3 (Ghaedi et al., 2000), acyl-CoA oxidase (Tsukamoto et al., 1990), catalase (Tsukamoto et al., 1990), ADAPS (Honsho et al., 2008), DHAPAT (Honsho et al., 2017), 3-ketoacyl-CoA thiolase (Tsukamoto et al., 1990), and guinea pig anti-Pex14 antibody (Mukai et al., 2002). Rabbit antiserum to phosphorylated Pex14 at Ser232, termed anti-Pex14-pS232 antibody, was raised in Biologica (Nagoya, Japan) by conventional subcutaneous injection of a synthetic 21-amino acid phosphopeptide comprising a 19-amino acid residues at 223-241 of rat Pex14 including a phospho-Ser232 and Gly-Cys di-peptide sequence at the C-terminus that had been linked to keyhole limpet hemocyanin (Tsukamoto et al., 1990). The raised rabbit antibody was purified in Biologica by affinity chromatography using a column conjugated to the synthetic phosphopeptide antigen after passing thorough that conjugated to the corresponding unmodified peptide. We purchased rabbit polyclonal antibodies to FLAG (Sigma-Aldrich) and Erk1/2 (Cell Signaling), mouse monoclonal antibodies to HA (16B12; Covance), FLAG (Sigma-Aldrich), hexa-histidine tag (Qiagen), GFP (Santa Cruz Biotechnology, Inc.), phospho-Erk1/2 (Cell Signaling), cytochrome P450 reductase (Santa Cruz Biotechnology, Inc.), cytochrome *c* (BD Pharmingen), β-actin (MBL), and Tom20 (Santa Cruz Biotechnology, Inc.), and goat anti-lactate dehydrogenase antibody (Rockland). Kinase inhibitors, U0126 and SB203580, were purchased from Cell Signaling. KU5933 and Compound C were from Abcam and Merck, respectively.

### Preparation of mouse tissues and Phos-tag PAGE

Several different tissues from an 8-week old male mouse that had been fed with normal chow under regular day-light and dark cycle were directly lysed in buffer-L (20 mM HEPES-KOH, pH 7.4, 0.15 M NaCl, 25 μg/ml each of leupeptin and antipain, 1 mM phenylmethylsulfonyl fluoride (PMSF), and 1 mM dithiothreitol) containing 0.5% Nonidet P-40 and 0.1% SDS by ten strokes of homogenization with an Elvehjem-Potter homogenizer (Miura et al., 1992). After centrifugation, solubilized fractions (15 µg) were subjected to SDS-PAGE as described (Natsuyama et al., 2013). In phosphatase treatment, the soluble fractions were incubated with 1 µg/ml of λ-protein phosphatase (New England Biolab) for 30 min at 30°C. Phos-tag PAGE was performed with 7.5% polyacrylamide gels containing 50 µM Phos-tag (Wako Chemicals) and 100 µM MnCl_2_ for the lysates of mouse tissues and 25 µM Phos-tag and 50 µM MnCl_2_ for those of cultured cells (Kinoshita et al., 2006).

### Mass spectrometry analysis

Fao cells (8 × 10^6^ cells) were lysed in RIPA buffer (50 mM Tris-HCl, pH7.6, 0.15 M NaCl, 1% Nonidet P-40, 0.5% sodium deoxycholate, 0.1% SDS, and 1 mM DTT supplemented with a complete protease inhibitor cocktail (Roche) and Phos-stop phosphatase inhibitor cocktail (Sigma-Aldrich) and were flash-frozen in liquid nitrogen. After thawing on ice and centrifugation at 20,000 *g* for 15 min at 4°C, supernatant fractions were subjected to immunoprecipitation with anti-Pex14 antibody immobilized on SureBeads Protein G magnetic beads (Bio-Rad) for 3 h at 4°C with rotation. After washing with RIPA buffer four times and then with 50 mM ammonium bicarbonate twice, proteins on the beads were digested by adding 400 ng trypsin/Lys-C mix (Promega) for 16 h at 37°C. The digests were acidified and desalted using GL-Tip SDB (GL Sciences). The eluates were evaporated in a SpeedVac concentrator and dissolved in 3% acetonitrile (ACN) and 0.1% trifluoroacetic acid.

LC-MS/MS analysis of the resultant peptides was performed on an EASY-nLC 1200 UHPLC connected to a Q Exactive Plus mass spectrometer equipped with a nanoelectrospray ion source (Thermo Fisher Scientific). The peptides were separated on a 75 µm inner diameter x 150 mm C18 reversed-phase column (Nikkyo Technos) with a linear 4**–**28% ACN gradient for 0**–**100 min followed by an increase to 80% ACN for 10 min. The mass spectrometer was operated in a data-dependent acquisition mode with a top 10 MS/MS method. MS1 spectra were measured with a resolution of 70,000, an automatic gain control (AGC) target of 1 × 10^6^ and a mass range from 350 to 1,500 *m/z*. HCD MS/MS spectra were acquired at a resolution of 17,500, an AGC target of 5 × 10^4^, an isolation window of 2.0 *m/z*, a maximum injection time of 60 ms and a normalized collision energy of 27. Dynamic exclusion was set to 10 sec. Raw data were directly analyzed against the SwissProt database restricted to *H*. *sapiens* using Proteome Discoverer version 2.3 (Thermo Fisher Scientific) with Mascot search engine version 2.5 (Matrix Science) for identification and label-free precursor ion quantification. The search parameters were as follows: (i) trypsin as an enzyme with up to two missed cleavages; (ii) precursor mass tolerance of 10 ppm; (iii) fragment mass tolerance of 0.02 Da; (iv) carbamidomethylation of cysteine as a fixed modification; and (v) acetylation of the protein N-terminus, oxidation of methionine and phosphorylation of serine, threonine, and tyrosine as variable modifications. Peptides were filtered at a false-discovery rate of 1% using the percolator node. Normalization was performed such that the total sum of abundance values for each sample over all peptides was the same.

### Immunofluorescence microscopy

Immunostaining of cells was performed as described (Okumoto et al., 2011) with 4% paraformaldehyde for cell fixation and 0.1% Triton X-100 for permeabilization. Immuno-complexes were visualized with Alexa Fluor 488-labeled goat anti-rabbit IgG antibody and Alexa Fluor 568-labeled goat anti-guinea pig IgG antibody (Invitrogen). Cells were observed by a confocal laser microscope (LSM710 with Axio Observer Z1; Zeiss) equipped with a Plan Apochromat 100x 1.4 NA oil immersion objective lens and argon plus dual HeNe lasers at RT. Images were acquired with Zen software (Zeiss) and prepared using Photoshop (CS4; Adobe).

### Subcellular fractionation and immunoprecipitation

For separation of cytosolic and organelle fractions from CHO cells, harvested cells were incubated with 25 μg/ml digitonin in buffer H (20 mM Hepes-KOH, pH 7.4, 0.25 M sucrose, 1 mM DTT, complete protease inhibitor cocktail [Roche], 1 mM NaF, 1 mM Na_3_VO_4_, and 6 mM β-glycerophosphate) for 5 min at room temperature as described (Natsuyama et al., 2013). After centrifugation at 20,000 *g* for 30 min at 4°C, equal aliquots of respective fractions were analyzed by immunoblotting. For isolation of organelle fraction of Fao cells, ∼4 × 10^6^ cells were homogenized with a Potter–Elvehjem teflon homogenizer (Wheaton) in buffer H and centrifuged at 800 *g* for 10 min at 4°C to yield post nuclear supernatant (PNS) fraction. Organelle fraction was separated by ultracentrifugation of PNS fraction at 100,000 *g* for 30 min at 4°C. The organelle pellet was lysed in buffer L containing 0.5% CHAPS, 1 mM NaF, 1 mM Na_3_VO_4_, and 6 mM β-glycerophosphate for 30 min at 4°C. After centrifugation at 20,000 *g* for 10 min at 4°C, resulting supernatants were incubated with anti-Pex14 antibody in buffer L containing 0.5% CHAPS for 2 h at 4°C. Antibody-antigen complexes were recovered by incubating for 1 h at 4°C with Protein A-Sepharose CL-4B (GE Healthcare) and eluted with Laemmli sample buffer. For detection of mono-ubiquitinated Pex5 in organelle fraction of Fao cells, subcellular fractionation was performed with H buffer containing 5 mM N-ethylmaleimide and no DTT as described (Okumoto et al., 2011).

### *In vitro* binding assay

Pex14, Pex5S, Pex5L, EGFP-PTS1, and HA-catalase were expressed as GST fusion proteins in *Escherichia coli* DH5α and were purified with glutathione-Sepharose beads (GE Healthcare), as described (Otera et al., 2002). Pex5S, Pex5L, EGFP-PTS1, and HA-catalase were isolated from the purified GST fusion proteins by cleaving with PreScission protease (GE Healthcare) according to the manufacturer’s protocol. GST or GST-Pex14 variants (typically 2 μg each) conjugated to glutathione-Sepharose beads were incubated with Pex5 (2 μg), Pex13 (0.1 μg), EGFP-PTS1 (4 μg), or HA-catalase (4 μg) by rotating for 2 h at 4°C in an *in vitro* binding buffer (50 mM Tris-HCl, pH 7.5, 0.15 M NaCl, 1% Triton X-100, 10% glycerol, 1 mM PMSF, 1 mM EDTA, and 1 mM DTT). Glutathione-Sepharose beads were washed three times with the binding assay buffer minus glycerol and the bound fractions were eluted with Laemmli sample buffer.

### Cell viability assay

Cell viability was measured with a tetrazolium-based toxicology assay kit (Promega). CHO cells (1 × 10^4^ cells per well) were seeded in 96-well plate and grown for 24 h and then treated for 14 h with 0.8 mM H_2_O_2_ alone or together with 20 mM 3-aminotriazole (3-AT). After treatment, cells were incubated with Celltiter 96 aqueous one solution reagent (Promega) for additional 2 h and the cell viability was determined by the absorbance of media at 490 nm as described in the manufacturer’s protocol.

### Pulse-chase experiment

Fao cells growing in DMEM supplemented with 10 % FBS in 6-well plate were washed twice with PBS, incubated in cysteine- and methionine-free DMEM (Gibco) supplemented with 10% FBS that had been dialyzed for 1 h in excess PBS with Slide-A-Lyzer dialysis cassette (Thermo Fisher Scientific). Cells were then pulse-labeled for 1 h by adding 100 μCi/ml ^35^S-methionine plus ^35^S-cysteine (American Radiolabeled Chemicals). To chase the ^35^S-labeled proteins, cells were washed twice and further incubated for 1 h with DMEM supplemented with 10 % FBS and 10 mM methionine. ^35^S-labeled cells were harvested and incubated for 5 min in buffer H containing 50 μg/ml digitonin at room temperature as described (Natsuyama et al., 2013). After centrifugation at 20,000 *g* for 30 min at 4°C, cytosolic and organelle fractions were subjected to immunoprecipitation with antibodies to catalase and DHAPAT as described (Tsukamoto et al., 1990). ^35^S-labeled proteins were separated by SDS-PAGE and detected with an Autoimaging analyzer (Typhoon FLA-9500; GE Healthcare).

### Statistical analysis

Statistical analysis was performed using R software (http://www.r-project.org). Quantitative data were represented as means ± SD from at least three independent experiments. Statistical significance was determined using a two-tailed unpaired Student’s *t* test for comparisons between two groups or one-way ANOVA with Dunnett’s post hoc test for more than two groups. P-values of <0.05 were considered statistically significant.

## Acknowledgments

We thank S. Okuno and Y. Nanri for technical assistance, the other members of Fujiki laboratory for expertise and discussions, and the members of Functional Cell Biology laboratory of Kyushu University for continuous supports. We thank Dr. Daniel Hess, Friedrich Miescher Institute for Biomedical Research, Basel, Switzerland, for LC/MS/MS analysis at the initial stage of this work. This work was supported in part by grants from the Ministry of Education, Culture, Sports, Science, and Technology of Japan; Grants-in-Aid for Scientific Research, MEXT KAKENHI Grant Number JP26116007 (to Y.F.) and the Japan Society for the Promotion of Science Grants-in-aid for Scientific Research, Japan Society for the Promotion of Science KAKENHI Grants Numbers JP24770130, JP26440032, and JP17K07310 (to K.O.) and JP24247038, JP25112518, JP25116717, JP15K14511, JP15K21743, and JP17H03675 (to Y.F.); grants from the Takeda Science Foundation (to Y.F.), the Naito Foundation (to Y.F.), the Japan Foundation for Applied Enzymology (to Y.F. and K.O.), the Novartis Foundation (Japan) for the Promotion of Science (to Y.F.), Joint Usage and Joint Research Programs of Institute of Advanced Medical Sciences, Tokushima University (to Y.F.), and Qdai-jump Research Program in Kyushu University (to K.O.).

## Author Contributions

Conceptualization, K.O., M.E.S., and Y.F.; Methodology, K.O., M.E.S., H.K., and Y.F.; Validation, K.O., H.K. and Y.F.; Formal Analysis, K.O., H.K., and R.N.; Investigation, K.O., M.E.S., M.N., H.K., R.N., and T.M.; Writing – Original Draft, K.O., H.K., and Y.F.; Writing – Review & Editing, K.O., H.K. and Y.F.; Visualization, K.O.; Supervision, Y.F.; Funding Acquisition, K.O. and Y.F.; Resources, Y.F.

## Supplemental information

**Supplementary Figure 1.**
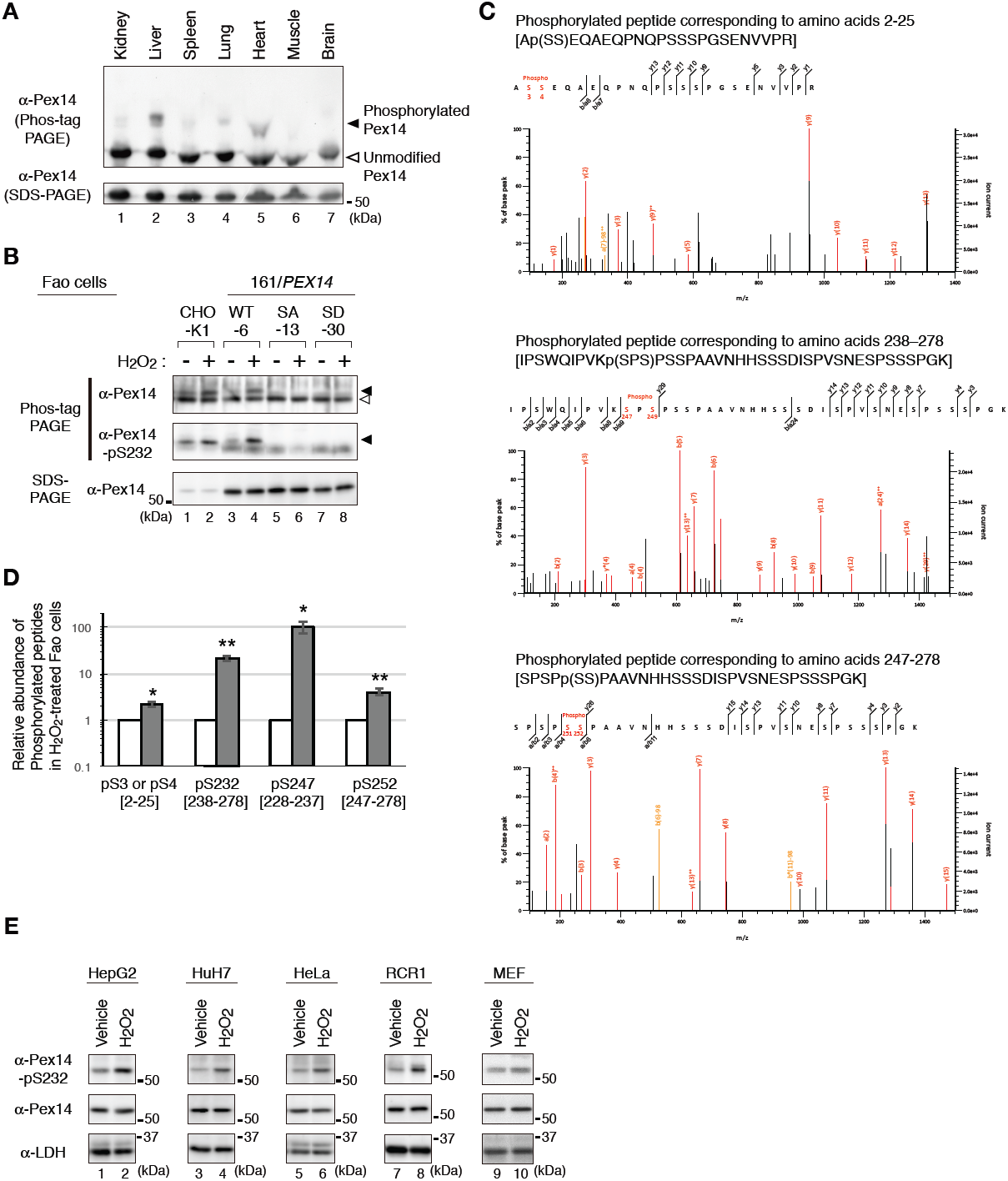
Pex14 is phosphorylated in mouse tissues and mammalian cultured cell lines. (A) Lysates of various mouse tissues (15 µg each) indicated at the top were analyzed by Phos-tag PAGE (upper panel), SDS-PAGE (lower panel), and immunoblotting with anti-Pex14 antibody. (B) CHO-K1 and *pex14* ZP161 stable cell lines each expressing wild-type His-Pex14 (WT-6), Pex14-S232A (SA-13), and Pex14-S232D (SD-30) were treated for 30 min with vehicle (-) and 0.8 mM H_2_O_2_. Cell lysates were analyzed by SDS-PAGE (upper panel), Phos-tag PAGE (lower panels), and immunoblotting with indicated antibodies. Open and solid arrowheads indicate unmodified and phosphorylated forms of His-Pex14, respectively. (C). LC-MS/MS analysis for phosphorylation sites of Pex14 upon H_2_O_2_ treatment was performed as in Fig. 2B. Fragment spectra of phosphorylated peptides corresponding to amino acids 2-25 containing pS3 or pS4 (upper panel), amino acids 238-278 containing pS247 (middle panel), and amino acids 247-278 containing pS252 (lower panel) are shown. (D) Quantification of phosphorylated Pex14 upon H_2_O_2_-treatment. Phosphorylated peptides were identified in Pex14 isolated from Fao cells that had been treated with vehicle or H_2_O_2_ as described in Fig. 2B. The levels of respective phosphopeptides in H_2_O_2_-treated cells (solid bars) were quantified with label-free precursor ion quantification and represented by taking as 1.0 that in vehicle-treated cells (open bars). Error bars represent means ± SEM of eight measurments in three independent experiments. *, *p* < 0.05; **, *p* < 0.01; unpaired Student’s *t* test versus vehicle treated cells. (E) Phosphorylation of Pex14 at Ser232 was induced in various cultured cells upon H_2_O_2_-treatment. Several mammalian cell lines (4 × 10^5^ cells each) were treated with vehicle and 0.5 mM H_2_O_2_ for 0.5 h (HepG2, HuH7, and MEF) and 1 h (HeLa and RCR1), similarly to Fig. 1B, and were analyzed by SDS-PAGE and immunoblotting with indicated antibodies.

**Supplementary Figure 2.**
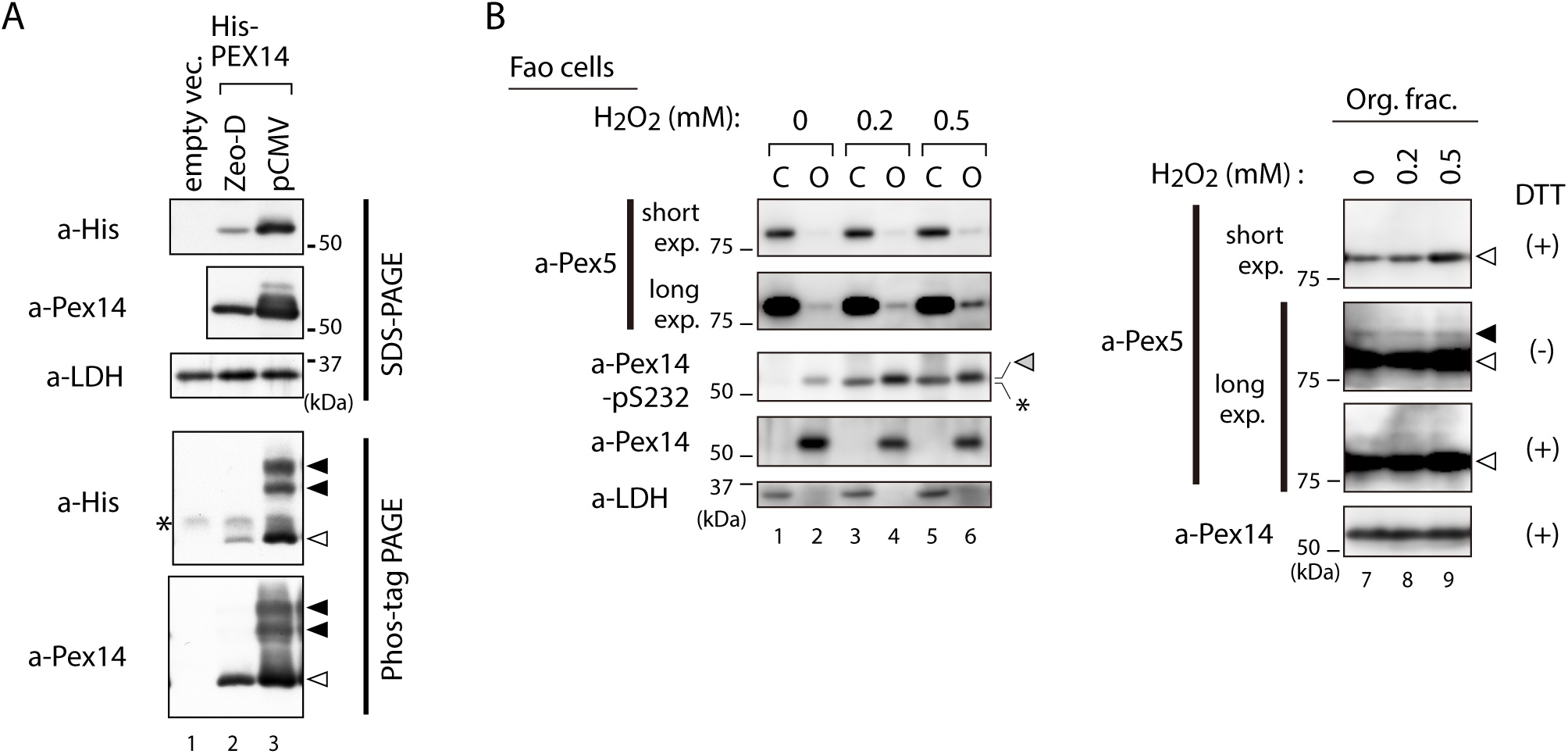
Effects of Pex14 phosphorylation on catalase import into peroxisomes. (A) Protein level of exogenously expressed Pex14. *pex14* ZP161 cells were transfected with a pCMVSPORT1 vector harboring no insert (mock, lane 1) or that encoding His-Pex14 (pCMV, lane 3), and pcDNAZeo-D vector encoding His-Pex14 (Zeo-D, lane 2) At 24-h after transfection, cells were lysed and analyzed by SDS-PAGE (upper panels) and Phos-tag PAGE (lower panels), and immunoblotting with indicated antibodies. Open and solid arrowheads indicate unmodified and phosphorylated forms of His-Pex14, respectively. *, a nonspecific band. (B) Pex5 recycling upon H_2_O_2_-treatment. Fao cells treated with H_2_O_2_ at indicated concentration for 1 h were homogenized in the presence 5 mM NEM. PNS fractions were separated into cytosolic (C) and organelle (O) fractions. Equal aliquots of the cytosolic and organelle fractions in Laemmli sample buffer with 0.1 M DTT (left panels) and organelle fractions in the presence (+) or absence (-) of DTT (right panels) were analyzed by SDS-PAGE and immunoblotting with indicated antibodies. Gray, solid, and open arrowheads indicate phosphorylated Pex14 at S232, mono-ubiquitinated Pex5 at Cys11 (Okumoto et al., 2011), and unmodified Pex5, respectively. *, a nonspecific band.

**Supplementary Figure 3.**
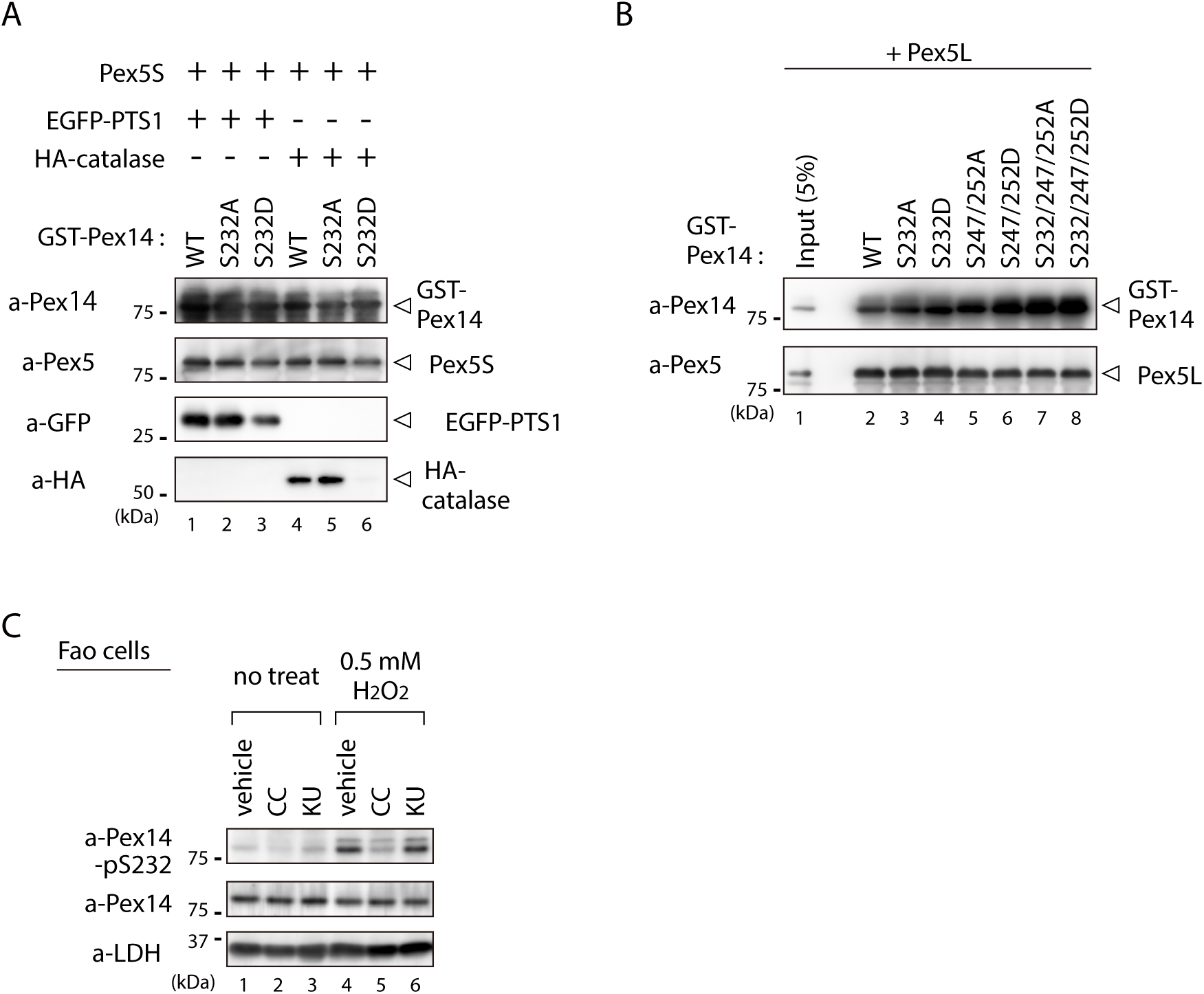
S232D mutation in Pex14 affects Pex5-catalase interaction. (A) Pex14-S232D also shows less affinity to catalase in the complex with Pex5S. *In vitro* binding assay was performed using GST-Pex14 variants, HA-catalase, and Pex5S as in Fig. 5B. (B) Direct interaction of Pex14 variants with Pex5L. GST pull-down assay was performed with a pair of GST-Pex14 variants with Pex5L as in Fig. 5B. (C) Effect of an ATM inhibitor KU-55933 on Pex14 phosphorylation. Fao cells pretreated for 30 min with DMSO (vehicle), 10 μM KU-55933 (KU), or 10 μM Compound C (CC)) were cultured for 1 h in the presence of 0.5 mM H_2_O_2_. Cell lysates were analyzed as in Fig. 1B by immunoblotting with the indicated antibodies.

## References

Abe, Y., Honsho, M., Itoh, R., Kawaguchi, R., Fujitani, M., Fujiwara, K., Hirokane, M., Matsuzaki, T., Nakayama, K., Ohgi, R., et al. (2018). Peroxisome biogenesis deficiency attenuates the BDNF-TrkB pathway-mediated development of the cerebellum. Life Sci. Alliance 1, e201800062.

Albuquerque, C.P., Smolka, M.B., Payne, S.H., Bafna, V., Eng, J., and Zhou, H. (2008). A multidimensional chromatography technology for in-depth phosphoproteome analysis. Mol. Cell. Proteomics 7, 1389–1396.

Apanasets, O., Grou, C.P., Van Veldhoven, P.P., Brees, C., Wang, B., Nordgren, M., Dodt, G., Azevedo, J.E., and Fransen, M. (2014). PEX5, the shuttling import receptor for peroxisomal matrix proteins, is a redox-sensitive protein. Traffic 15, 94–103.

Carvalho, A.F., Pinto, M.P., Grou, C.P., Alencastre, I.S., Fransen, M., Sá-Miranda, C., and Azevedo, J.E. (2007). Ubiquitination of mammalian Pex5p, the peroxisomal import receptor. J. Biol. Chem. 282, 31267–31272.

de Vet, E.C.J.M., IJlst, L., Oostheim, W., Wanders, R.J.A., and van den Bosch, H. (1998). Alkyl-dihydroxyacetonephosphate synthase. Fate in peroxisome biogenesis disorders and identification of the point mutation underlying a single enzyme deficiency J. Biol. Chem. 273, 10296–10301.

Farre, J.-C., Manjithaya, R., Mathewson, R.D., and Subramani, S. (2008). PpAtg30 tags peroxisomes for turnover by selective autophagy. Dev. Cell 14, 365–376.

Fransen, M., Nordgren, M., Wang, B., and Apanasets, O. (2012). Role of peroxisomes in ROS/RNS-metabolism: Implications for human disease. Biochim. Biophys. Acta-Mol. Basis Dis. 1822, 1363–1373.

Fujiki, Y., Miyata, N., Mukai, S., Okumoto, K., and Cheng, E.H. (2017). BAK regulates catalase release from peroxisomes. Mol. Cell. Oncol. 4, article: e1306610.

Fujiki, Y., Okumoto, K., Mukai, S., Honsho, M., and Tamura, S. (2014). Peroxisome biogenesis in mammalian cells. Front. Physiol. 5, article 307.

Gaestel, M. (2006). MAPKAP kinases - MKs - two’s company, three’s a crowd. Nat. Rev. Mol. Cell Biol. 7, 120–130.

Gerbeth, C., Schmidt, O., Rao, S., Harbauer, A.B., Mikropoulou, D., Opalinska, M., Guiard, B., Pfanner, N., and Meisinger, C. (2013). Glucose-induced regulation of protein import receptor Tom22 by cytosolic and mitochondria-bound kinases. Cell Metab. 18, 578–587.

Ghaedi, K., Tamura, S., Okumoto, K., Matsuzono, Y., and Fujiki, Y. (2000). The peroxin Pex3p initiates membrane assembly in peroxisome biogenesis. Mol. Biol. Cell 11, 2085–2102.

Gould, S.J., Keller, G.-A., and Subramani, S. (1987). Identification of a peroxisomal targeting signal at the carboxy terminus of firefly luciferase. J. Cell Biol. 105, 2923–2931.

Hartman, P., Belmont, P., Zuber, S., Ishii, N., and Anderson, J. (2003). Relationship between catalase and life span in recombinant inbred strains of *Caenorhabditis elegans*. J. Nematol. 35, 314–319.

Honsho, M., Abe, Y., and Fujiki, Y. (2017). Plasmalogen biosynthesis is spatiotemporally regulated by sensing plasmalogens in the inner leaflet of plasma membranes. Sci. Rep. 7, 43936.

Honsho, M., Yagita, Y., Kinoshita, N., and Fujiki, Y. (2008). Isolation and characterization of mutant animal cell line defective in alkyl-dihydroxyacetonephosphate synthase: Localization and transport of plasmalogens to post-Golgi compartments. Biochim. Biophys. Acta 1783, 1857–1865.

Hosoi, K., Miyata, N., Mukai, S., Furuki, S., Okumoto, K., Cheng, E.H., and Fujiki, Y. (2017). The VDAC2–BAK axis regulates peroxisomal membrane permeability. J. Cell Biol. 216, 709–721.

Hwang, I., Lee, J., Huh, J.Y., Park, J., Lee, H.B., Ho, Y.S., and Ha, H. (2012). Catalase deficiency accelerates diabetic renal injury through peroxisomal dysfunction. Diabetes 61, 728–738.

Itoh, R., and Fujiki, Y. (2006). Functional domains and dynamic assembly of the peroxin Pex14p, the entry site of matrix proteins. J. Biol. Chem. 281, 10196–10205.

Ivashchenko, O., Van Veldhoven, P.P., Brees, C., Ho, Y.S., Terlecky, S.R., and Fransen, M. (2011). Intraperoxisomal redox balance in mammalian cells: oxidative stress and interorganellar cross-talk. Mol. Biol. Cell 22, 1440–1451.

Johnson, M.A., Snyder, W.B., Cereghino, J.L., Veenhuis, M., Subramani, S., and Cregg, J.M. (2001). *Pichia pastoris* Pex14p, a phosphorylated peroxisomal membrane protein, is part of a PTS-receptor docking complex and interacts with many peroxins. Yeast 18, 621–641.

Kinoshita, E., Kinoshita-Kikuta, E., Takiyama, K., and Koike, T. (2006). Phosphate-binding tag, a new tool to visualize phosphorylated proteins. Mol. Cell Proteomics 5, 749–757.

Koepke, J.I., Nakrieko, K.A., Wood, C.S., Boucher, K.K., Terlecky, L.J., Walton, P.A., and Terlecky, S.R. (2007). Restoration of peroxisomal catalase import in a model of human cellular aging. Traffic 8, 1590–1600.

Komori, M., Kiel, J.A.K.W., and Veenhuis, M. (1999). The peroxisomal membrane protein Pex14p of *Hansenula polymorpha* is phosphorylated in vivo. FEBS Lett. 457, 397–399.

Legakis, J.E., Koepke, J.I., Jedeszko, C., Barlaskar, F., Terlecky, L.J., Edwards, H.J., Walton, P.A., and Terlecky, S.R. (2002). Peroxisome senescence in human fibroblasts. Mol. Biol. Cell 13, 4243–4255.

Liu, X., Ma, C., and Subramani, S. (2012). Recent advances in peroxisomal matrix protein import. Curr. Opin. Cell Biol. 24, 1–6.

Meinecke, M., Cizmowski, C., Schliebs, W., Krüger, V., Beck, S. R. W., and Erdmann, R. (2010). The peroxisomal importomer constitutes a large and highly dynamic pore. Nat. Cell Biol. 12, 273–277.

Middelkoop, E., Wiemer, E.A., Schoenmaker, D.E., Strijland, A., and Tager, J.M. (1993). Topology of catalase assembly in human skin fibroblasts. Biochim. Biophys. Acta 1220, 15–20.

Miura, S., Kasuya-Arai, I., Mori, H., Miyazawa, S., Osumi, T., Hashimoto, T., and Fujiki, Y. (1992). Carboxyl-terminal consensus Ser-Lys-Leu-related tripeptide of peroxisomal proteins functions *in vitro* as a minimal peroxisome-targeting signal. J. Biol. Chem. 267, 14405–14411.

Miyazawa, S., Osumi, T., Hashimoto, T., Ohno, K., Miura, S., and Fujiki, Y. (1989). Peroxisome targeting signal of rat liver acyl-coenzyme A oxidase resides at the carboxy terminus. Mol. Cell. Biol. 9, 83–91.

Mukai, S., and Fujiki, Y. (2006). Molecular mechanisms of import of peroxisome-targeting signal type 2 (PTS2) proteins by PTS2 receptor Pex7p and PTS1 receptor Pex5pL. J. Biol. Chem. 281, 37311–37320.

Mukai, S., Ghaedi, K., and Fujiki, Y. (2002). Intracellular localization, function, and dysfunction of the peroxisome-targeting signal type 2 receptor, Pex7p, in mammalian cells. J. Biol. Chem. 277, 9548–9561.

Natsuyama, R., Okumoto, K., and Fujiki, Y. (2013). Pex5p stabilizes Pex14p: a study using a newly isolated *pex5* CHO cell mutant, ZPEG101. Biochem. J. 449, 195–207.

Oeljeklaus, S., Schummer, A., Mastalski, T., Platta, H.W., and Warscheid, B. (2016). Regulation of peroxisome dynamics by phosphorylation. Biochim. Biophys. Acta. 1863, 1027–1037.

Okatsu, K., Oka, T., Iguchi, M., Imamura, K., Kosako, H., Tani, N., Kimura, M., Go, E., Koyano, F., Funayama, M., et al. (2012). PINK1 autophosphorylation upon membrane potential dissipation is essential for Parkin recruitment to damaged mitochondria. Nat. Commun. 3, 1016.

Okumoto, K., Abe, I., and Fujiki, Y. (2000). Molecular anatomy of the peroxin Pex12p: RING finger domain is essential for Pex12p function and interacts with the peroxisome-targeting signal type 1-receptor Pex5p and a RING peroxin, Pex10p. J. Biol. Chem. 275, 25700–25710.

Okumoto, K., Misono, S., Miyata, N., Matsumoto, Y., Mukai, S., and Fujiki, Y. (2011). Cysteine ubiquitination of PTS1 receptor Pex5p regulates Pex5p recycling. Traffic 12, 1067–1083.

Osumi, T., Tsukamoto, T., Hata, S., Yokota, S., Miura, S., Fujiki, Y., Hijikata, M., Miyazawa, S., and Hashimoto, T. (1991). Amino-terminal presequence of the precursor of peroxisomal 3-ketoacyl-CoA thiolase is a cleavable signal peptide for peroxisomal targeting. Biochem. Biophys. Res. Commun. 181, 947–954.

Otera, H., and Fujiki, Y. (2012). Pex5p imports folded tetrameric catalase by interaction with Pex13p. Trafffic 13, 1364–1377.

Otera, H., Harano, T., Honsho, M., Ghaedi, K., Mukai, S., Tanaka, A., Kawai, A., Shimizu, N., and Fujiki, Y. (2000). The mammalian peroxin Pex5pL, the longer isoform of the mobile peroxisome targeting signal (PTS) type 1 transporter, translocates Pex7p-PTS2 protein complex into peroxisomes via its initial docking site, Pex14p. J. Biol. Chem. 275, 21703–21714.

Otera, H., Okumoto, K., Tateishi, K., Ikoma, Y., Matsuda, E., Nishimura, M., Tsukamoto, T., Osumi, T., Ohashi, K., Higuchi, O., et al. (1998). Peroxisome targeting signal type 1 (PTS1) receptor is involved in import of both PTS1 and PTS2: Studies with *PEX5*-defective CHO cell mutants. Mol. Cell. Biol. 18, 388–399.

Otera, H., Setoguchi, K., Hamasaki, M., Kumashiro, T., Shimizu, N., and Fujiki, Y. (2002). Peroxisomal targeting signal receptor Pex5p interacts with cargoes and import machinery components in a spatiotemporally differentiated manner: conserved Pex5p WXXXF/Y motifs are critical for matrix protein import. Mol. Cell. Biol. 22, 1639–1655.

Platta, H.W., Brinkmeier, R., Reidick, C., Galiani, S., Clausen, M.P., and Eggeling, C. (2016). Regulation of peroxisomal matrix protein import by ubiquitination. Biochim. Biophys. Acta - Mol. Cell Res. 1863, 838–849.

Purdue, P.E., and Lazarow, P.B. (1996). Targeting of human catalase to peroxisomes is dependent upon a novel COOH-terminal peroxisomal targeting sequence. J. Cell Biol. 134, 849–862.

Ray, P.D., Huang, B.W., and Tsuji, Y. (2012). Reactive oxygen species (ROS) homeostasis and redox regulation in cellular signaling. Cell Signal. 24, 981–990.

Schliebs, W., Saidowsky, J., Angianian, B., Dodt, G., Herberg, F.W., and Kunau, W.-H. (1999). Recombinant human peroxisomal targeting signal receptor PEX5. Structural basis for interaction of PEX5 with PEX14. J. Biol. Chem. 274, 5666–5673.

Schmidt, O., Harbauer, A.B., Rao, S., Eyrich, B., Zahedi, R.P., Stojanovski, D., Schoenfisch, B., Guiard, B., Sickmann, A., Pfanner, N., et al. (2011). Regulation of mitochondrial protein import by cytosolic kinases. Cell 144, 227–239.

Schrader, M., and Fahimi, H.D. (2006). Peroxisomes and oxidative stress. Biochim. Biophys. Acta-Mol. Cell Res. 1763, 1755–1766.

Shimizu, N., Itoh, R., Hirono, Y., Otera, H., Ghaedi, K., Tateishi, K., Tamura, S., Okumoto, K., Harano, T., Mukai, S., et al. (1999). The peroxin Pex14p: cDNA cloning by functional complementation on a Chinese hamster ovary cell mutant, characterization, and functional analysis. J. Biol. Chem. 274, 12593–12604.

Sies, H., Berndt, C., and Jones, D.P. (2017). Oxidative Stress. Annu. Rev. Biochem. 86, 715–748.

Swinkels, B.W., Gould, S.J., Bodnar, A.G., Rachubinski, R.A., and Subramani, S. (1991). A novel, cleavable peroxisomal targeting signal at the amino-terminus of the rat 3-ketoacyl-CoA thiolase. EMBO J. 10, 3255–3262.

Tada-Oikawa, S., Hiraku, Y., Kawanishi, M., and Kawanishi, S. (2003). Mechanism for generation of hydrogen peroxide and change of mitochondrial membrane potential during rotenone-induced apoptosis. Life Sci. 73, 3277–3288.

Tanaka, K., Soeda, M., Hashimoto, Y., Takenaka, S., and Komori, M. (2013). Identification of phosphorylation sites in *Hansenula polymorpha* Pex14p by mass spectrometry. FEBS Open Bio. 3, 6–10.

Tsukamoto, T., Yokota, S., and Fujiki, Y. (1990). Isolation and characterization of Chinese hamster ovary cell mutants defective in assembly of peroxisomes. J. Cell Biol. 110, 651–660.

Walton, P.A., Brees, C., Lismont, C., Apanasets, O., and Fransen, M. (2017). The peroxisomal import receptor PEX5 functions as a stress sensor, retaining catalase in the cytosol in times of oxidative stress. Biochim Biophys Acta-Mol. Cell Res. 1864, 1833–1843.

Waterham, H.R., Ferdinandusse, S., and Wanders, R.J.A. (2016). Human disorders of peroxisome metabolism and biogenesis. Biochim. Biophys. Acta-Mol. Cell Res. 1863, 922–933.

Weiner, M.P., Costa, G.L., Schoettlin, W., Cline, J., Mathur, E., and Bauer, J.C. (1994). Site-directed mutagenesis of double-stranded DNA by the polymerase chain reaction. Gene 151, 119–123.

Will, G.K., Soukupova, M., Hong, X., Erdmann, K.S., Kiel, J.A.K.W., Dodt, G., Kunau, W.-H., and Erdmann, R. (1999). Identification and characterization of the human orthologue of yeast Pex14p. Mol. Cell Biol. 19, 2265–2277.

Ye, G., Metreveli, N.S., Donthi, R.V., Xia, S., Xu, M., Carlson, E.C., and Epstein, P.N. (2004). Catalase protects cardiomyocyte function in models of type 1 and type 2 diabetes. Diabetes 53, 1336–1343.

Zhang, J., Tripathi, D.N., Jing, J., Alexander, A., Kim, J., Powell, R.T., Dere, R., Tait-Mulder, J., Lee, J.-H., Paull, T.T., et al. (2015). ATM functions at the peroxisome to induce pexophagy in response to ROS. Nat. Cell Biol. 17, 1259–1269.

